# Microbial communities in an ultra-oligotrophic sea are more affected by season than by distance from shore

**DOI:** 10.1101/2020.04.17.044305

**Authors:** Markus Haber, Dalit Roth Rosenberg, Maya Lalzar, Ilia Burgsdorf, Kumar Saurav, Regina Lionheart, Yoav Lehahn, Dikla Aharonovich, Daniel Sher, Michael D. Krom, Laura Steindler

**Affiliations:** Department of Marine Biology, Leon H. Charney School of Marine Sciences, University of Haifa, Israel; Department of Aquatic Microbial Ecology, Institute of Hydrobiology, Biology Centre CAS, Czech Republic; Bioinformatics Service Unit, University of Haifa, Israel; The Dr. Moses Strauss Department of Marine Geosciences, Leon H. Charney School of Marine Sciences, University of Haifa, Israel; Morris Kahn Marine Research Station, Environmental Geochemistry Lab., Leon H. Charney School of Marine Sciences, University of Haifa, Israel

**Keywords:** Mediterranean Sea, SAR11, Transect, Seasonality, 16S rRNA

## Abstract

Marine microbial communities vary seasonally and spatially, but these two factors are rarely addressed together. We studied temporal and spatial patterns of the microbial community structure and activity along a coast to offshore transect from the Israeli coast of the Eastern Mediterranean Sea (EMS) over six cruises, in three seasons of two consecutive years. The ultra-oligotrophic status of the South Eastern Mediterranean Sea was reflected in the microbial community composition that was dominated by oligotrophic microbial groups such as SAR11 throughout the year, even at the most coastal station sampled. Seasons affected microbial communities much more than distance from shore explaining about half of the observed variability in the microbial community, compared to only about 6% that was explained by station. However, the most coastal site differed significantly in community structure and activity from the three further offshore stations in early winter and summer, but not in spring. Our data on the microbial community composition and its seasonality from a transect into the South Eastern Levantine basin support the notion that the EMS behaves similar to open gyres rather than an inland sea.

## Introduction

Marine microbial communities play a pivotal function in the biogeochemistry of the ocean because of their key roles in the carbon, nitrogen and sulfur cycles (Falkowski et al. 2008). Their composition is strongly affected by seasons with recurring microbial turnover over different years as revealed by oceanographic time series studies conducted both at offshore stations (*e.g.* Bermuda Atlantic Time-Series Study (BATS) in the Western Atlantic Ocean and the Hawaii Ocean Time-series (HOT) in the North Pacific subtropical gyre (Giovannoni and Vergin 2012)) and more coastal affected sites (*e.g.* the San Pedro Ocean Time Series (SPOTS) in southern California (Fuhrman et al. 2006) and the western English Channel (Gilbert et al. 2012)). Spatial variability of marine surface water microbial communities has been reported at various scales (Ghiglione et al. 2005; Sunagawa et al. 2015). The observed differences were linked to gradients in environmental conditions (Fortunato et al. 2011; Sunagawa et al. 2015; Wang et al. 2019) and distinct water masses with different physico-chemical properties (Dubinsky et al. 2017; Morales et al. 2018). Spatial variability due to environmental gradients is especially evident when comparing nearshore and offshore microbial communities as parameters such as nutrient availability, temperature, dissolved organic matter, and, especially at estuaries, salinity change from the coast to the open ocean (Ghiglione et al. 2005; Fortunato et al. 2011; Quero and Luna 2014; Lucas et al. 2016; Wang et al. 2019). These parameters are influenced by season leading to weakening or strengthening of the environmental gradients along coast to offshore transects. Few studies have followed seasonal differences in the microbial community dynamics over a coastal to offshore transect (Fortunato et al. 2011; Lucas et al. 2016; Wang et al. 2019), so relatively little is known about how these temporal and spatial changes interact and affect the microbial community composition and their functional potential. Only one of these studies (Wang et al. 2019) addressed community structure changes without the confounding factor of salinity, which has been identified as a key factor in structuring microbial communities at estuaries (Fortunato et al. 2011).

Here we followed the seasonal dynamics of microbial communities along a coast to offshore transect in the ultra-oligotrophic Eastern Mediterranean Sea (EMS). This mostly land-enclosed region, represents the largest body of water that is severely depleted in phosphate (Krom et al. 1991), and one of the most oligotrophic oceanic regions on Earth (Krom et al. 2010). Despite being an inland sea with major anthropogenic external nutrient supply, it has been suggested that it behaves similar to open ocean gyres as a result of its unusual anti-estuarine circulation (Powley et al. 2017). The nutrient cycle at offshore sites in the EMS follows predictable patterns. During summer the water column is well stratified with a distinct deep chlorophyll maximum (DCM) and very low nutrient content in the surface waters (Kress and Herut 2001; Krom et al. 2014). In winter, as a result of increasingly deep-water mixing, dissolved nutrients are advected into the photic zone. Due to local weather conditions (short cold and wet (often stormy) periods interspersed with clear sunny ones), the phytoplankton bloom in this region starts soon after the nutrients are supplied to the photic zone and increases throughout the winter to a maximum in late winter (Krom et al. 2014). The dissolved phosphate in surface waters is consumed during the winter phytoplankton bloom, while measurable nitrate persists (Krom et al. 1992). In the summer autotrophs in offshore waters of this region tend to be phosphate and nitrogen co-limited, while heterotrophic bacteria are either phosphate or phosphate and nitrogen co-limited (Thingstad et al. 2005; Tanaka et al. 2011; Tsiola et al. 2016). Detailed molecular characterization of the microbial community in this geographic area are limited to few studies that represent only snapshots of the microbial communities taken at a single time point (Feingersch et al. 2010; Keuter et al. 2015; Dubinsky et al. 2017). Until now molecular studies investigating seasonal and spatial differences in the microbial community structure focused on nitrogen fixing microbes using clone libraries of the *nifH* (Man-Aharonovich et al. 2007; Yogev et al. 2011). However, nitrogen fixation seems rare in the euphotic zone of the Eastern Mediterranean (Yogev et al. 2011) and thus these microbes are not expected to reflect a large part of the microbial community. The goal of the present study was to examine both microbial community structure and activity in time and space by amplicon sequencing of 16S rRNA genes and transcripts. Surface seawater samples (10 m) were collected along a coastal-to-offshore transect in three seasons (early winter, spring and summer) and for two consecutive years. The oceanographic status of the system was determined from physical, chemical and remote sensing measurements, and the phytoplankton community analyzed by flow cytometry and pigment analysis.

## Material and Methods

### Sampling

Six one-day cruises were performed onboard the R/V Mediterranean Explorer over a two-year period: two in early winter (December 1^st^, 2014; November 17^th^, 2015), two in spring (March 24^th^, 2015; March 30^th^, 2016) and two in summer (July 14^th^, 2015; July 25^th^, 2016). Cruises followed the same coastal to offshore transect heading out from the Herzliya marina, Israel (Figure 1). Four stations, labeled station 1 to 4 in coastal to offshore order, were analyzed for different *in situ* parameters using a conductivity, temperature and depth (CTD) probe. Seawater was collected from 10 m depth using twelve 8L Niskin bottles mounted on a rosette for nutrient, pigment, cell count, DNA and RNA analyses. To better identify gradients along the transect, CTD and nutrient data were collected for two additional stations: one located between stations 1 and 2, and the other between station 2 and 3. Station coordinates, bottom depths, distance between stations and to the shore are summarized in Supplementary Table S1.

**Figure 1.**
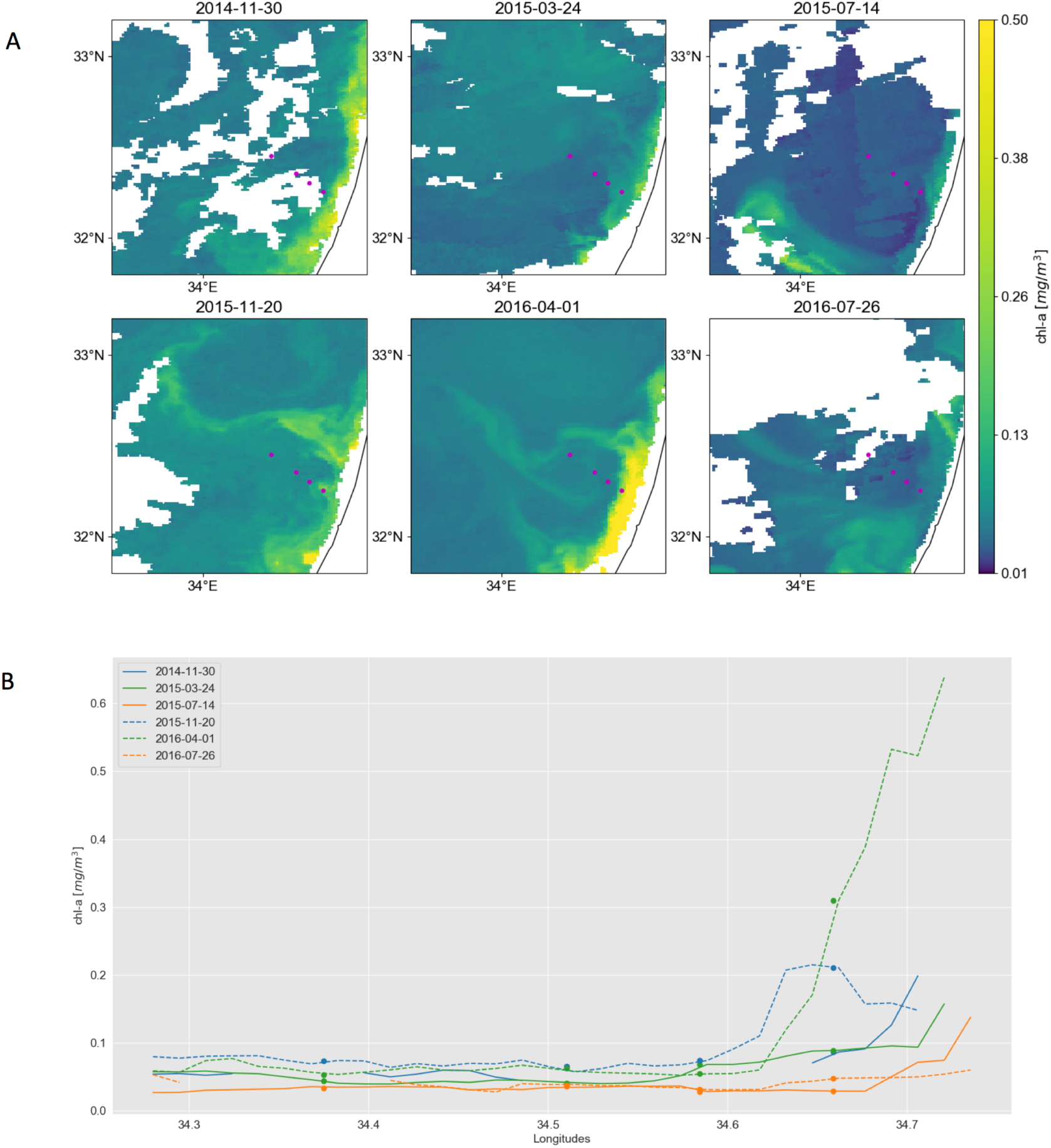
Surface chlorophyll concentrations in the southeastern Mediterranean. Dots mark locations of sampling stations, which are from right (coast) to left offshore: St1, St2, St3, St4. A) Maps derived from Copernicus Marine Environment Monitoring Service (CMEMS) merged satellite data. To reduce the area masked by clouds (white color), chlorophyll maps from 30.11.2014, 20.11.2015, 01.04.2016 and 26.07.2016 were used instead of 01.12.2014, 17.11.2015, 30.03.2016 and 25.07.2016, respectively. Furthermore, maps for 30.11.2014, 24.3.2015, 14.7.2015 and 26.7.2016 are composed of 5, 3, 3 and 3 consecutive images, respectively. Purple dots mark the locations of sampling stations. B) Surface chlorophyll concentrations along a coast-to-open-sea transect overlapping the sampling stations. The transects are derived from the multi-satellite chlorophyll maps shown in A, with each line corresponding to a different sampling date.

### CTD data and surface chlorophyll *a* maps

Satellite-based maps of surface chlorophyll concentration are derived from the Copernicus Marine Environment Monitoring Service (CMEMS, http://marine.copernicus.eu/services-portfolio/access-toproducts/). We used the level-3 Mediterranean Sea reprocessed surface chlorophyll concentration (OCEANCOLOUR_MED_CHL_L3_REP_OBSERVATIONS_009_073) product, which consists of merged SeaWiFS, MODIS-Aqua, MERIS and VIIRS satellite data. Using multi-satellite data allows continuous tracking of fine-scale chlorophyll filaments as they are advected and deformed by the currents (Lehahn et al. 2017). Surface chlorophyll concentration are estimated via the MedOC4 (Volpe et al. 2007) and the AD4 (D’Alimonte and Zibordi 2003) algorithms for case-1 and case-2 waters, respectively. Spatial and temporal resolution is 1 km and 1 day, respectively.

A SeaBird CTD profiler (SBE 19plus V2) was used to measure vertical profiles of temperature and salinity up to a depth of 500 m. Chlorophyll *a* fluorescence profiles were measured with a Seapoint fluorometer calibrated with bottle chlorophyll measurements and turbidity profiles with a Seapoint turbidity sensor, both mounted on the CTD. Data were extracted using the Seasoft V2 software suite and plotted using Ocean Data View 4.7.4 (Schlitzer R. 2015, http://odv.awi.de).

### Nutrient analysis

Seawater for nutrient analysis was collected in 15 ml Falcon tubes pre-rinsed with sample seawater. For each nutrient, duplicate non-filtered samples were frozen onboard directly after collection and kept at −20°C until analysis of silicate, nitrate+nitrite and soluble reactive phosphate content (within six weeks) at the service unit of the Interuniversity Institute for Marine Sciences in Eilat, Israel. During the July 2016 cruise the same nutrients plus ammonia were also measured in non-frozen samples. These samples were filtered through 0.4 µm filters, transferred into 15 ml Falcon tubes, stored at 4°C and analysed within 24 hours at the Morris Kahn Marine Laboratory, Sdot Yam, Israel using a SEAL AA-3 autoanalyzer. Methods are described in the Supplementary method information.

### Flow cytometry

For flow cytometry sample collection, 1.5 ml triplicate seawater samples were fixed with glutaraldehyde (0.125% final concentration), incubated in the dark for 10 min, stored in liquid nitrogen onboard and kept at −80 °C in the lab until analysis. Then, samples were thawed at room temperature and run on a BD FACSCanto™ II Flow Cytometry Analyzer Systems (BD Biosciences) for counts of phytoplankton and total cell counts and an easyCyte HT Guava flow cytometer (Merck Millipore) for total cell counts (see Supplementary method information for details).

### Pigment analysis

Phytoplankton community structure was identified by pigment analysis. Four to eleven L seawater were filtered on Glass fiber filters (25 mm GF/F, Whatman, nominal pore size 0.7µm). Filters were dried by placing their underside on a kimwipe, transferred to cryovials, stored in liquid nitrogen until arrival at the laboratory, and kept at −80°C until extraction. The collected cells were extracted in 1 ml 100% methanol for 2.5 h at room temperature. Extractions were immediately clarified with syringe filters (Acrodisc CR, 13 mm, 0.2 µm PTFE membranes, Pall Life Sciences) and transferred to UPLC vials. Samples were run on an UPLC and pigments identified based on retention time and spectrum absorbance. Several known standards (DHI Water and Environment Institute, (Hørsholm, Denmark)) were used to ease identification and calculate pigment concentrations (see Supplementary Information for details).

### DNA and RNA sample collection and extraction

Seawater was collected from the Niskin bottles into 10 or 20 L polycarbonate carboys. Tubes for water transfer and the carboys were pre-rinsed three times with sample water. Samples for DNA (5 to 11.5 L) were pre-filtered through 11 and 5 µm nylon filters (Millipore), cells collected on 0.22 µm sterivex filters (Millipore) and kept in storage buffer (40 mM EDTA, 50 mM Tris pH 8.3, 0.75 M sucrose). Samples for RNA (0.5 to 4.5 L, according to volume filtered within 15 minutes from Niskin bottles being on-deck) were collected without pre-filtration, directly on 0.2 µm filters (Supor-200 Membrane Disc Filters, 25 mm; Pall Corporation) and filters preserved in RNA Save (Biological Industries). DNA and RNA samples were stored in liquid nitrogen onboard and kept at −80°C in the lab until extraction. Nucleic acids were extracted at the BioRap unit, Faculty of Medicine, Technion, Israel using a semi-automated protocol, which includes manually performed chemical and mechanical cell lysis before the automated steps (see Supplementary Information for details).

### PCR amplification and sequencing of 16S rRNA genes (DNA) and transcripts (RNA) samples

Prior to reverse transcription, all RNA samples were tested by PCR for the presence of contaminating DNA. Total RNA was reverse transcribed using the iScript cDNA synthesis kit (BioRad) according to the manufacturer’s instructions. A two-stage “targeted amplicon sequencing” protocol (*e.g.* (Green et al. 2015)) was performed to PCR amplify the 16S rRNA gene from cDNA and DNA (see Supplementary Information for a detailed description). The primers used in the first PCR stage consisted of the 16S primer set 515F-Y and 926R (Parada et al. 2016) that targets the variable V4-5 region with common sequence tags added at the 5’ end as described previously (*e.g.* (Moonsamy et al. 2013)). The first PCR stage was performed in triplicates, which were pooled after validation on 1% agarose gels. Subsequently, a second PCR amplification was performed to prepare libraries. These were pooled and after a quality control sequenced (2×250 paired end reads) using an Illumina MiSeq sequencer. Library preparation and pooling were performed at the DNA Services (DNAS) facility, Research Resources Center (RRC), University of Illinois at Chicago (UIC). MiSeq sequencing was performed at the W.M. Keck Center for Comparative and Functional Genomics at the University of Illinois at Urbana-Champaign (UIUC).

### Sequence processing

After quality control of the obtained pair of fastq files for each sample, the paired reads were assembled and additional QC steps taken with MOTHUR Version 1.40.2 (Schloss et al. 2009) (see Supplementary Information for details). The filtered dataset was clustered into operational taxonomic units (OTUs) at 97% similarity using QIIME. Representative sequences for each OTU were obtained and classified using MOTHUR with the silva v128 database at 80% confidence level. OTUs identified as non-prokaryotic, chloroplasts- or mitochondria-origin and OTUs with <10 reads across all samples were removed. OTUs identified as SAR11 were also classified against Silva v132 to get clade assignment. In total, the 56 samples had 2,820,858 quality sequences, which binned into 9448 OTUs. In order to avoid bias related to differences in library size, all libraries were rarified to 20,000 reads per sample using the R package Vegan (Oksanen et al. 2017). 9396 OTUs based on 97% similarity were retained after subsampling. Of these, 8372 OTUs were found in DNA samples with 1538 to 2253 OTUs present per sample and 8684 OTUs in RNA samples ranging from 1604 to 2230 OTUs per sample. Read data were deposited in the NCBI SRA database under the project number PRJNA548664. Station 4 data are labeled N1200, a change made to ease reading of the article.

### Sequencing controls and variance among replicates

Four negative controls from the first PCR were added randomly onto the sequencing plates to monitor overall potential for cross contaminations in both PCR and sequencing. These negative control samples averaged 929 reads, compared to samples averaging 50,503 reads. Accordingly, contaminant DNA should poorly compete with sample DNA for amplification.

To assess within station variability and robustness of the observed trends, duplicates for DNA and RNA samples were collected and extracted for samples from station 4 for the summer 2015 cruise, the spring 2015 and 2016 cruises, as well as for station 1 for the spring 2016 cruise. We chose these samples to have a representation from different seasons, as well as from the most coastal and most offshore stations. Based on their Bray-Curtis dissimilarity, replicates were significant closer to each other than to other stations from the same cruise (paired Wilcoxon signed-rank test, RNA: z=3.4044 and *P*<0.001; DNA: z=4.2812 and *P*<0.001).

### Microbial community analyses

Alpha diversity parameters (Chao1 richness, Shannon H’ diversity and Simpson index of dominance) were calculated using the R package iNEXT (Chao et al. 2014; Hsieh et al. 2018). Beta-diversity was estimated by pairwise calculation of the Bray-Curtis dissimilarity with the R package Vegan and used for non-metric multidimensional scaling analysis. Vegan was also used to examine significance of grouping factors (such as molecule type, season and sampling station) with the ADONIS test and to examine correspondence between abiotic and biotic measurements and variation in community composition by variation partitioning analysis and canonical correspondence analysis, for which proposed explanatory variables were divided into three matrices: I) Physical: distance from shore, temperature, salinity, turbidity; II) Nutrients: phosphate, nitrate+nitrite, silicate; III) Biological: total fluorescence, total cell counts, *Prochlorococcus* cell counts, *Synechococcus* cell counts (see Supplementary method information for details of all Vegan analyses).

For differential OTU abundance analysis between sample groups (*e.g.* molecule type, season, sampling station), OTU count data was normalized using the cumulative sum scaling method implemented in the R package metagenomeSeq (Paulson et al. 2013) and analyzed with the R package DEseq2 (Love et al. 2014). Significant differential OTU abundance was based on Benjamini-Hochberg adjusted *P*-values.

Synchronous dynamics of single OTUs were analyzed by soft clustering with the R package Mfuzz (Kumar and Futschik 2007) using OTUs that comprised more than 0.1% of total reads of DNA and RNA samples, respectively. OTU tables were standardized using the normal SD based method, fuzzifier variables were estimated (DNA: 1.19569, RNA: 1.182411) and cluster number was set to 10. Only OTUs with cluster membership values of at least 0.7 were considered.

Similarity profile analysis (SIMPROF) (Clarke et al. 2008) based on Bray-Curtis similarity was run in R using the clustsig package version 1.1 to test for structure in the spring 2016 samples relating to the intrusion of coastal waters into offshore waters.

### Statistical analyses

Paired Wilcoxon signed rank tests were performed to test if Bray-Curtis dissimilarity between replicates from the same sample location was lower than between samples from different locations from the same cruise. A Kruskal-Wallis test was used to find significant difference in alpha diversity indices between seasons. Two-tailed Mann-Whitney U tests were used to determine for each season if Bray-Curtis dissimilarity between station 1 and the other three station was higher than between the three further offshore station. Finally, Spearman correlations were performed to compare the relative abundance in RNA and flow cytometry data for *Synechococcus* and *Prochlorococcus*, to compare if distance between stations correlates with Bray-Curtis dissimilarity between samples and to compare if differences in relative abundance in RNA and DNA data are correlated. All these tests were performed in PAST version 3.14 (Hammer et al. 2001). Other statistical tests were performed as described above and implemented in the used R packages.

## Results

### Environmental setting

The previously reported ultraoligotrophic status of the Levantine basin (Krom et al. 2005) was reflected in our nutrient data of samples collected at 10 m (Supplementary Table S3 and S4). Phosphate concentrations obtained from frozen samples were near or below our detection limit of 40 nM in all cruises with a maximum of 70 nM at station 1 in early winter 2014.

Nitrate+nitrite concentrations for these samples ranged from < 50 nM to 240 nM at the most offshore station and from <50 to 690 nM at the most coastal station. They were below our detection limit (50 nM) for frozen samples during summer cruises apart from station 1 in the July 2015 cruise (120 nM). In early winter they decreased from station 1 towards the offshore stations, whereas in spring they increased from station 1 to station 3 before decreasing again towards station 4, the most offshore station. In the summer 2016 cruise, we performed also an analysis on fresh (unfrozen) samples that provided a lower limit of detection (Krom et al. 2005). This enabled the identification of the nutrient range during the most nutrient depleted state.

Concentrations of phosphate ranged from 0.2 to 2.1 nM, of nitrate from 8 to 28 nM, and of ammonia from 1.3 to 7 nM (Supplementary Table S4) again confirming the nutrient depleted status of the Levantine Basin during summer.

Temperature and salinity varied seasonally at the sampling depth (10 m). The lowest temperatures were observed in spring (17.7-18.6°C) and the highest in summer (26.2-28.7°C). Seasonal changes in salinity were small (<0.7 PSU from lowest to highest value) (Supplementary Table S2).

Together CTD (Supplementary Figure 1), nutrient (Supplementary Table S3) and satellite data (Figure 1) indicated that our sampling times corresponded to three distinct stages of the annual cycle: the stratified water column in summer, early mixing and phytoplankton bloom in early winter and the declining phytoplankton bloom in spring (Krom et al. 1992; Kress and Herut 2001). Satellite data indicated the peak of the phytoplankton bloom to be in both years in February (data not shown). Based on pigment data, early winter samples were dominated by haptophytes of either Phaeocystaceae or of members of the Prymnesiaceae and Isochrysidaceae families (see details on pigment results in Supplementary information as well as Supplementary Figure S2 and Supplementary Table S5). Spring samples were characterized by declining phytoplankton bloom communities (Supplementary Figure S2, Supplementary Table S5) and peak abundances of *Prochlorococcus* (Supplementary Figure S5B and Table S6). During summer the water column was stratified with a mixed layer depth of 30m and characterized by nutrient depleted surface waters (Supplementary Table S2) and the lowest *Synechococcus* abundances at stations 2 to 4 (Supplementary Figure S5A, Supplementary Table S6).

When comparing the transect sampling stations from the coast outwards, a clear difference was seen between station 1, the shallowest and most coastal sampling site, and the other three more offshore stations. Station 1 often differed from all other sampled stations in nutrient concentrations (*e.g.* silicate (in spring) and nitrite+nitrate (early winter, Supplementary Tables S2 and S3)), in remotely sensed surface chlorophyll (Figure 1) and several *in situ* parameters such as temperature (in early winter and spring), salinity (in summer), fluorescence (in early winter) and turbidity (in early winter and summer, Supplementary Table S5, Supplementary Figure S1). In accordance with (Berman et al. 1986), it was thus regarded as a coastal station.

### Microbial community composition

93 of the 9396 OTUs (based on 97% similarity) formed the “core” of the microbial community and were present in all samples and represented 61% of the total reads. Bacteria dominated with 99.68% of the total reads compared to 0.32% of reads identified as archaea. 95.3% of all reads belonged to nine bacterial classes: Alphaproteobacteria (48.2% of all reads), Cyanobacteria (17.6%), Gammaproteobacteria (15.4%), Flavobacteriia (5.4%), Marinimicrobia (SAR406 clade) (2.4%), the Verrucomicrobia class Opitutae (2.1%), Deltaproteobacteria (1.6%), Acidimicrobiia (1.3%) and Betaproteobacteria (1.3%). Most Alphaproteobacteria reads belonged to the SAR11 clade (60.4% of the Alphaproteobacteria), most reads of the class Cyanobacteria were identified as either *Prochlorococcus* or *Synechococcus* (together 85.9% of the Cyanobacteria), and the SAR86 clade was most common within the Gammaproteobacteria (32.8% of the Gammaproteobacteria). Relative abundances of the 13 families with >1% of total reads are shown in Figure 2.

**Figure 2.**
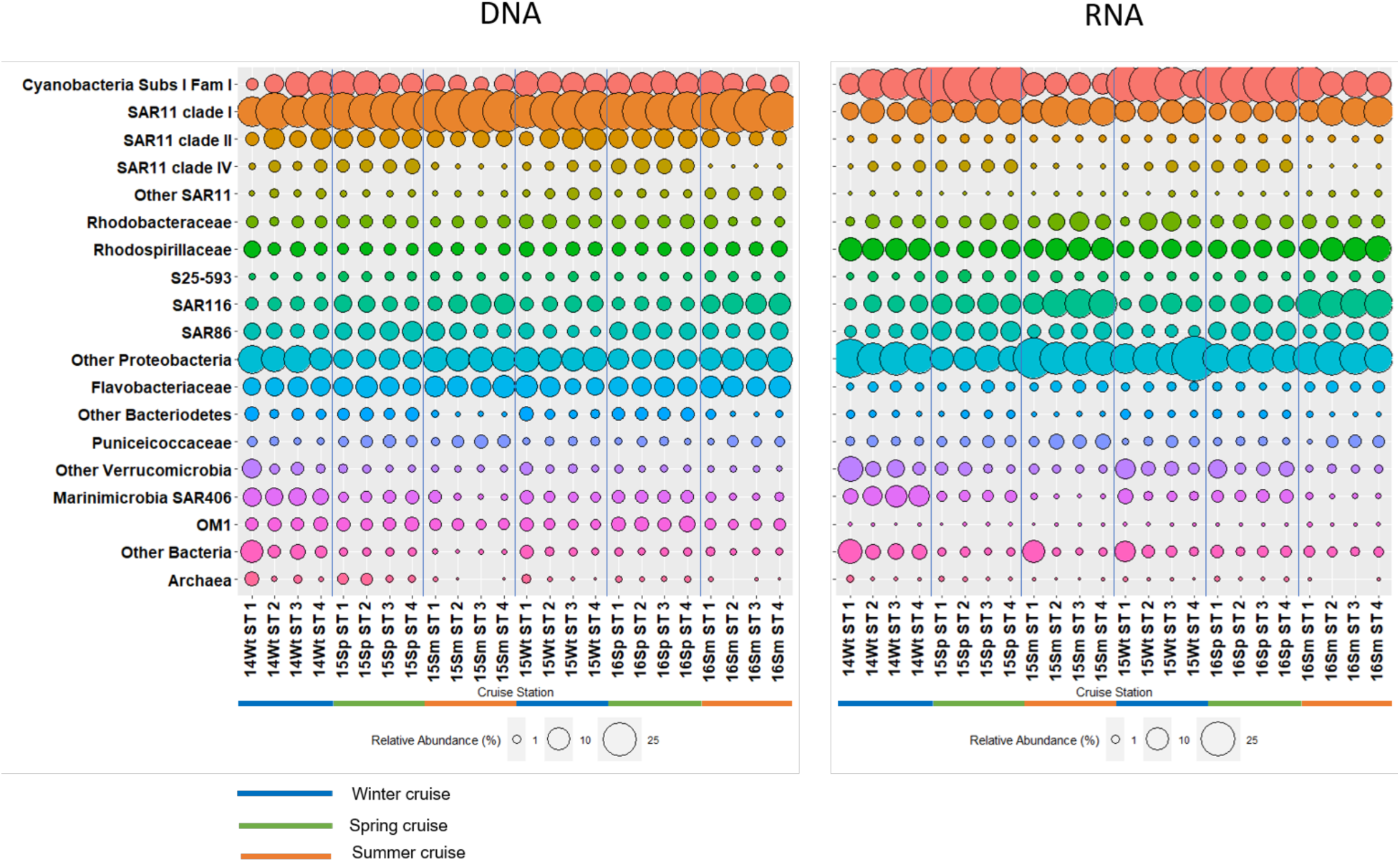
Relative abundance of microbial taxa in DNA (left) and RNA (right) samples. Families with > 1% total reads (DNA+RNA) are shown. Families below this threshold are summarized at the phylum level, except SAR11, which are summarized as other SAR11. Phyla with < 1% of total reads are summarized at the domain level. Color bars under the graphs indicate seasons: winter- blue, spring – green, summer – orange. Sample code: Last two digits of collection year (20XX), season (Wt – early winter, Sp - spring, Sm - summer) and station. Blue vertical lines separate different cruises.

A significant positive correlation (Spearman correlation, r>0, *P*<0.05) between relative abundance in DNA and RNA samples was present in 10 of these 13 families (Supplementary Table S7) indicating a strong link between presence and activity. However, the composition of resident (DNA) and active (RNA) microbial communities clearly differed (Figure 2) as supported by non-metric multidimensional scaling analysis (NMDS) (Figure 3) and Adonis test (F=22.37, R^2^=0.293, *P*<0.001). DEseq2 analysis identified 976 OTUs (representing 74.5% of all reads) with significantly different relative abundances between DNA and RNA (Benjamini- Hochberg adjusted *P*<0.05). Of these, 586 OTUs were more abundant in RNA samples and 390 in DNA samples. All significant OTUs from SAR11 clade I, clade II, the OM1 clade and the Flavobacteriaceae were higher in the DNA than the RNA samples. Significant OTUs of the Rhodobacteraceae, the SAR86 clade, the SAR116 and the Cyanobacteria subsection I family I, which included *Cyanobium*, *Synechococcus*, *Prochlorococcus* and OTUs unclassified at the genus level, were all higher in the RNA than in the DNA samples. Rhodospirillaceae and Marinimicrobia had both more OTUs with higher relative abundances in RNA than in DNA samples, however a few OTUs from both groups also showed the opposite trend. The remaining 309 significant OTUs did not sum up to >1% of total reads in any family. Given the difference between DNA and RNA samples, all subsequent analyses of microbial community composition (DNA) and activity (RNA) were performed separately.

**Figure 3.**
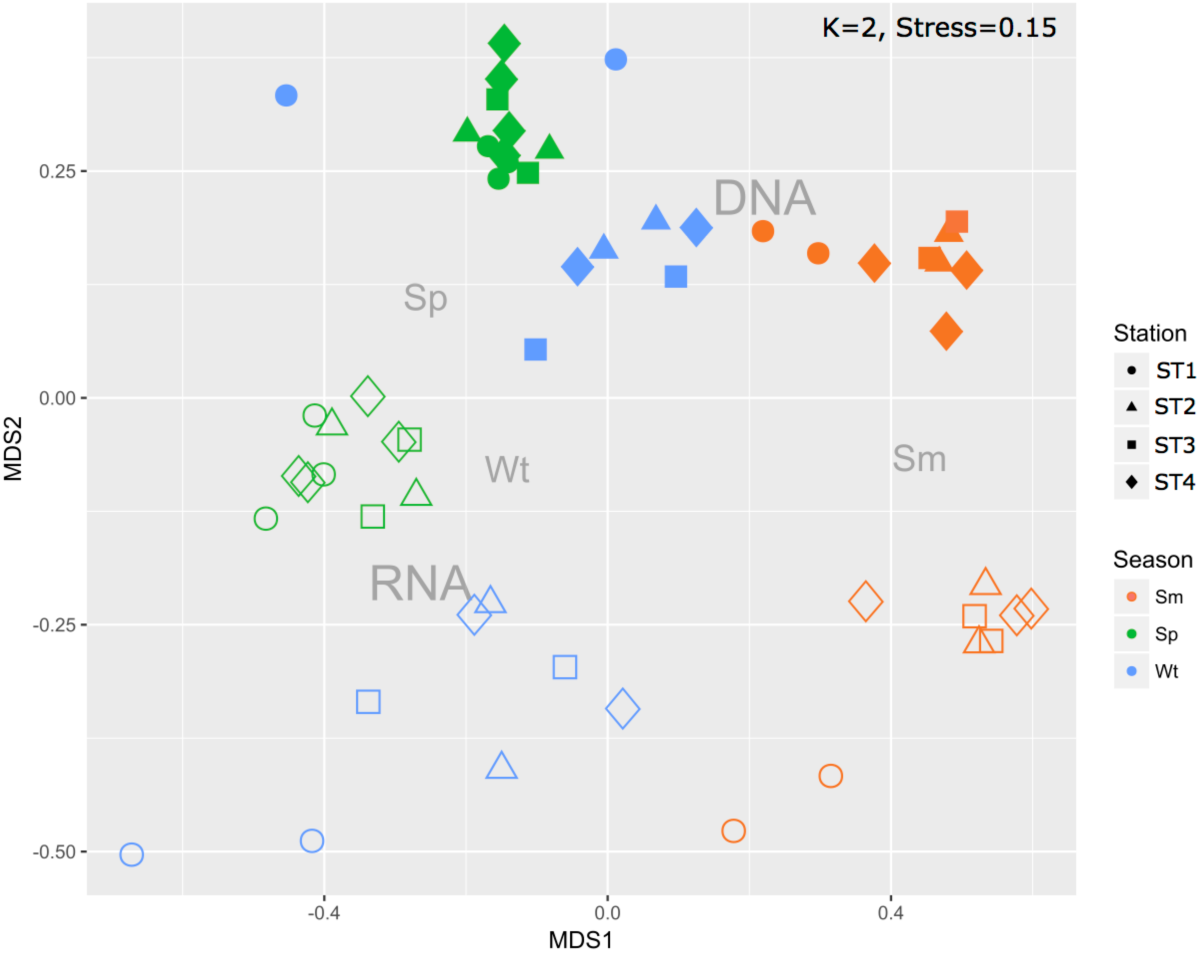
NMDS plot of the RNA (empty shapes) and DNA (filled shapes) samples. Colors represent seasons: Spring – green, Summer – red, early Winter – blue.

### Season had a larger effect on microbial structure and activity than spatial location

Season clearly affected both the microbial community structure (DNA) and activity (RNA) (Figure 3). This seasonal effect was much larger than the effect of station location across the coastal to offshore transect (DNA season R^2^=0.4980, station R^2^=0.0584; RNA season R^2^=0.5332, station R^2^=0.0607) with a small, significant interaction between season and station (DNA: R^2^=0.0840; RNA: R^2^=0.0806) (Adonis test, details in Supplementary Table S8).

Season affected also the microbial diversity. For DNA samples the three alpha diversity indices indicated lowest diversity in summer (Kruskal-Wallis test, *P*<0.001), when the system was most oligotrophic. The same pattern was observed for RNA samples, except for one index (Simpson diversity) that showed lowest diversity in spring (Supplementary Figure S3).

Soft clustering analysis on abundant OTUs was performed to identify OTUs that show the same abundance pattern across different seasons. Considering only OTUs with more than 0.1% of total reads, 98 of 110 OTUs at the DNA level, and 115 of 131 OTUs at the RNA level, were successfully clustered. At both the DNA and RNA level 8 of 10 clusters showed seasonal preferences (Figure 4). The OTUs of these seasonal clusters contributed 63.9% and 65.0% of total DNA and RNA reads, respectively. OTUs present in both clustering analyses showed consistent seasonal preferences (Supplementary Table S9). No OTU was assigned to more than one cluster. Cluster assignment, taxonomic identity and individual preferences of OTUs as well as abundance patterns of the non-seasonal clusters are shown in Supplementary File S1.

**Figure 4.**
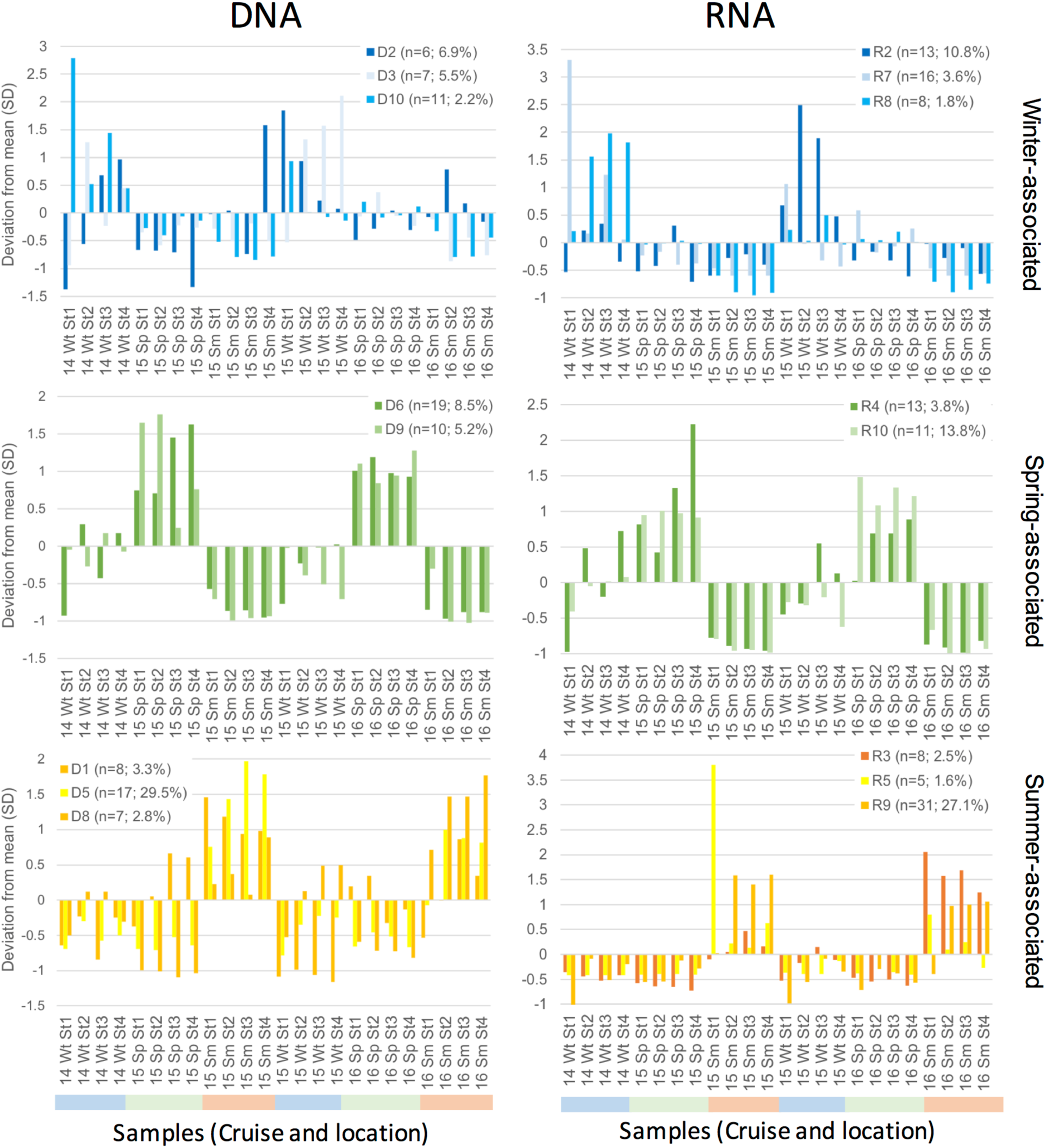
Seasonal patterns of OTUs with >0.1% of total reads in DNA and RNA based on soft cluster analysis. Numbers in brackets indicate number of OTUs in the cluster and the sum of their relative abundance in total DNA and RNA reads, respectively. Average relative abundance was set to 0 and y axis depicts change in abundance in standard deviations. X axis give samples sorted by cruise and location. Sample code consists of last two digits of collection year, season and station (St1 most coastal, St4 most offshore). Supplementary File S1 shows the patterns of the non-seasonal clusters D4, D7, R1 and R6 and data of OTUs that did not cluster.

DEseq2 analysis further confirmed that many OTUs differed significantly between at least two seasons. 582 OTUs representing 71.3% of all reads at the DNA level and 947 OTUs representing 81% of all reads at the RNA level had significant differences between two seasons (Benjamini- Hochberg adjusted *P*-value <0.05). Differences, in terms of relative abundance and number of significantly differing OTUs, were largest when comparing summer with the other two seasons (Supplementary File S2).

DEseq2 and soft clustering results identified several taxonomic groups favoring specific seasons: *e.g.* the Chloroflexi clade SAR202, the Marinimicrobia clade SAR406 and the Deltaproteobacteria clade SAR324 were more abundant in early winter, while the Aegan-169 marine group of the Alphaproteobacteria family Rhodospirillaceae were more abundant in summer. Lower species richness in summer, as indicated by lower Chao1 indices, was already evident at the order level (Supplementary Figure S3C). The majority of orders had no OTU that preferred summer over either spring or winter (Supplementary File S2).

At the DNA level, SAR11 was the order most affected by season, with seasonal preference varying according to SAR11 clade (Supplementary Figure 4). SAR11 clade Ia was the main SAR11 clade (51.4- 80.3% of all SAR11 DNA reads) and the most abundant taxonomic group overall (15.7-40.8% of the DNA reads). Its highest abundances were observed in summer.

Cluster analysis grouped its four most abundant OTUs (contributing 0.5 to 20.7% of total DNA reads) with summer-associated clusters, while the other clustered OTUs were assigned to a spring-associated cluster (Supplementary File 1). SAR11 clade Ia was the only SAR11 clade with summer associated OTUs in the cluster analysis apart from SAR11 clade III, which had one OTU but overall did not show a clear seasonal pattern. SAR11 clades Ib, II and IV had their lowest relative abundances in summer. SAR11 clade II peaked in spring and early winter, while clade IV was most abundant in spring (Supplementary Figure S4 B). Overall, few SAR11 OTUs differed significantly between winter and spring with all but one clade III OTU preferring spring (Supplementary file 2).

At the RNA level, cyanobacteria subsection I was the order that was mostly affected by season. All significant OTUs in the DEseq2 analysis (Benjamini-Hochberg adjusted *P*-value < 0.05) (125 OTUs, 22.6% of all RNA reads) belonged to family I, with most (83) being taxonomically assigned to *Prochlorococcus*. The vast majority of significant OTUs (109 OTUs) were significantly more active in spring than summer and 47 were significantly more active in early winter than summer. 43 OTUs differed significantly between spring and early winter (34 more active in spring, 9 in early winter). The few OTUs that were significantly more active in summer were all very rare. Overall cyanobacteria subsection I represented a larger part of the active community in spring (35.8% average of RNA reads), followed by winter (22.4%) and summer (12.9%) (Figure 2 B). *Prochlorococcus* made up for a larger part of the active community in spring, whereas no clear preference between winter and spring was observed for *Synechoccocus*. Their activity pattern correlated well with their cell abundances as determined by flow cytometry (Spearman rs 0.75 and 0.62, both *p*<0.001 for *Prochlorococcus* and *Synechococcus*, respectively) (Supplementary Figure S5).

Other taxonomic groups had closely related OTUs with different seasonal preferences. Details on their seasonal preferences are given in Supplementary Files S1 and S2, that show the cluster-based preferences of all abundant (>0.1 % of reads) OTUs at DNA and RNA levels and all OTUs with significant seasonal difference according to DEseq2 analysis (Benjamini-Hochberg adjusted *P*-value < 0.05), respectively.

### Spatial effects: Microbial communities at coastal station 1 differ from those at offshore stations in summer and early winter, but not in spring

Sampling site had a weak, yet significant effect on microbial structure and activity and interacted with season (Adonis test, see Supplementary Table S8 for details). However, no significant correlation was found between distance between stations and Bray-Curtis dissimilarity in microbial community structure (DNA) or activity (RNA) (Spearman correlation, *P*>0.05) irrespective if the whole data set was analyzed or each season separately. Further, when testing which environmental factors affected microbial community structure, distance between station was not a significant factor in the DNA analysis (canonical correspondence analysis (CCA), *P*>0.05, Supplementary Figure S7). In the RNA analysis distance was a significant factor in the CCA, but it was no longer significant when correcting for variance explained by the biological and nutrient data matrices (conditional CCA, *P*>0.05, Supplementary Figure S7).

As station 1 differed in environmental parameters from the three more offshore stations (see environmental settings above), we tested if this difference could be detected in the microbial community structure (DNA) and activity (RNA). Bray-Curtis dissimilarities were significantly higher between station 1 and the other three stations than among the three offshore stations, in both early winter and summer (Mann-Whitney U test, 2-tailed, winter: RNA z=-2.3219 *P*=0.020; DNA: z=-2.8022 *P*=0.005; summer: RNA z=-3.1825 *P*=0.001; DNA: z=-2.8353 *P*=0.005), but not in spring (RNA: z=-0.2967 *P*=0.767; DNA: z=-1.2858 *P*=0.199) (Figure 5). The effect was also observed in the NMDS plot as summer and early winter samples from station 1 grouped apart from the samples of stations 2 to 4 (Figure 3).

**Figure 5.**
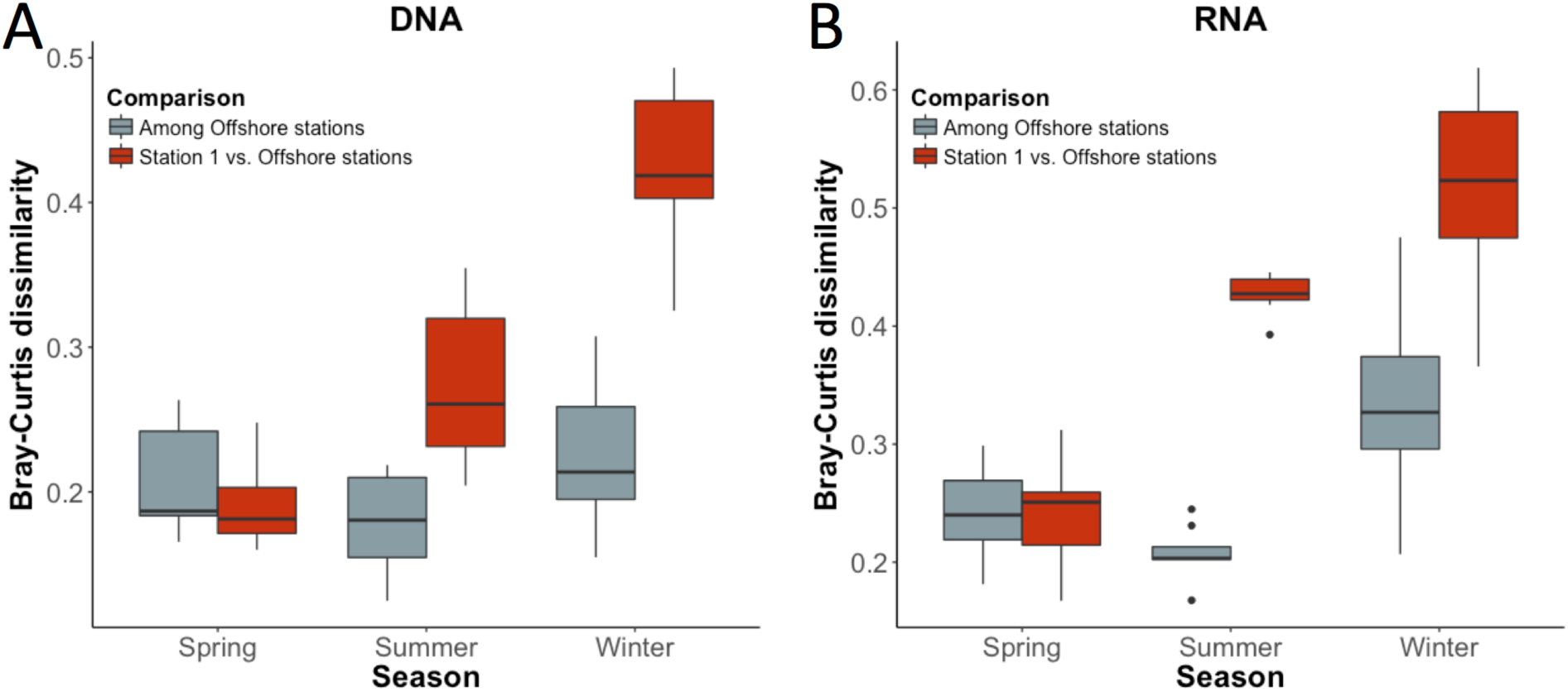
Bray-Curtis dissimilarity of A) DNA and B) RNA samples. Comparisons involving station 1 differ significantly from comparisons among stations 2, 3 and 4 in summer and early winter in both RNA and DNA samples

In summer, 22 OTUs (5.7% of all summer reads) and 27 OTUs (8.5% of all summer reads) at the DNA and RNA level, respectively, differed significantly between station 1 and stations 2-4 (DEseq2 analysis, Benjamini-Hochberg adjusted *P*-value < 0.05). In early winter 54 OTUs (5.0% of all early winter reads) and 76 OTUs (11.8% of all early winter reads) at the DNA and RNA level respectively, differed significantly between the coastal station and the offshore stations (DEseq2 analysis, Benjamini-Hochberg adjusted *P*-value < 0.05). 5 OTUs at the DNA level and 2 OTUs at the RNA differed significantly between station 1 and stations 2-4 in both summer and winter. Their preference was the same in both seasons suggesting a general preference for coastal or offshore conditions. Supplementary File S3 provides information on the taxonomy of groups enriched at station 1 or at the offshore stations 2-4.

### Offshore microbial communities are affected by intrusions of coastal water

At sites close to the Israeli coast of the EMS, short-term natural pulses of nutrients in the ultra-oligotrophic offshore waters can occur due to intrusions of mesoscale patches of coastal water that can be frequently observed by analysis of satellite-derived surface chlorophyll data (Efrati et al. 2013; Dubinsky et al. 2017). These intrusions of coastal water lead to shifts in the microbial communities of offshore affected sites (Dubinsky et al. 2017). Satellite chlorophyll (Figure 1) as well as *in situ* data (Supplementary Table S5) indicated an intrusion of coastal water into offshore waters taking place during the spring 2016 cruise, that affected station 4. As a result, phytoplankton community structure changed, as evidenced by increased concentrations in several pigments (Supplementary Table S4) and slightly higher abundances in *Synechococcus* cell numbers at the coastal station 1 and the intrusion-affected offshore station 4 compared to the offshore non-affected stations 2 and 3 (Supplementary Table S6). The affected stations grouped together in hierarchical clustering of the microbial structure (DNA) and activity (RNA) (Supplementary Figure S6), but DEseq2 analysis failed to identify any OTUs responsible for this difference (Benjamini-Hochberg adjusted *P*-value > 0.05 for all OTUs).

### Effects of environmental parameters on the microbial community structure and activity

Three data matrixes of external factors were examined for their influence on the microbial structure and activity: i) physical data (consisting of distance from shore, temperature, salinity and turbidity), ii) biological data: fluorescence (as proxy for chlorophyll), total cell number, *Prochlorococcus* counts, *Synechococcus* counts and iii) nutrient data: concentrations of phosphate, nitrate+nitrite, silicate. Together the three matrixes explained 49% of the observed variability of the microbial structure (DNA samples) and 72% of the microbial activity (RNA samples). Partition analysis indicated that in both analyses, the physical data matrix explained most of the variability (34.2% and 54.2% for DNA and RNA, respectively), followed by the biological data matrix (29.2% and 45.1% for DNA and RNA, respectively) and the nutrient data matrix (11.8% and 10.3% for DNA and RNA, respectively). The physical data matrix also had the largest proportion of variability not explained by any of the other two matrixes (Supplementary Figure S7).

A CCA analysis was performed to identify the environmental factors likely responsible for the observed patterns. Similar to the partition analysis, the physical matrix explained the largest amount of variance, followed by the biological matrix and the nutrient matrix. Temperature and salinity from the physical matrix and fluorescence from the biological matrix were significant factors in the DNA and RNA analysis (both in the unconditioned and conditioned analysis, *P*<0.05, Supplementary Figure S7). In the nutrient matrix, only silicate was consistently significant, but only in the DNA analysis (*P*<0.05, Supplementary Figure S7).

## Discussion

The ultra-oligotrophic nature of the Eastern Mediterranean Sea (EMS) is reflected in the microbial community structure of this coastal-offshore transect. Typical oligotrophic groups (*e.g.* SAR11) dominated throughout the year confirming previous snapshot studies of the EMS (Feingersch et al. 2010; Dubinsky et al. 2017). The microbial communities resembled closely those found at ultra-oligotrophic open ocean stations, both at the coastal station (*e.g.* SAR11 ranging from 26 ± 4 % in winter to 44 ± 6 % in summer) and even more at the offshore stations (*e.g.* SAR11 ranging from 44 ± 8 % in winter to 47 ± 5 % in summer). These estimates are similar if not higher than those found at oligotrophic ocean gyres such as the South Atlantic gyre (36 ± 9%) (Morris et al. 2012), the North Pacific gyre (ALOHA station, 44% in winter, 33% in summer) (Eiler et al. 2009), the South Pacific gyre (up to 53% in the most oligotrophic station) (West et al. 2016) and the Sargasso sea (North Atlantic gyre, BATS station, 33 ± 8%) (Carlson et al. 2009). Powley et al. (Powley et al. 2017) recently proposed that the EMS behaves similar to open gyres with respect to the importance of external nutrient supply, of dissolved organic matter as a key source of nutrients and ultra-oligotrophic conditions. Here we show that this is true also in terms of microbial community composition. The dominance of microbial groups typically considered to be oligotrophic at the coastal station (station 1) during spring and summer underlines that in the EMS oligotrophic conditions are found also close to the coast. SAR11 relative abundances were in fact much higher (>40% of DNA reads in both spring and summer) than those found at coastal stations in other parts of the Mediterranean Sea, including the North Western Mediterranean and the Adriatic Sea (Alonso-Sáez et al. 2007; Quero and Luna 2014; Tinta et al. 2015), highlighting the overall more oligotrophic status of the South Eastern Levantine basin.

In our study, season affected the microbial community structure and activity much more than spatial location along the coastal to offshore transect. This finding differs from other studies that analyzed spatial-temporal community changes in coastal-offshore transects. In a transect that sampled the Columbia river, its estuary, plume and coastal to offshore lines, Fortunato et al (Fortunato et al. 2011) described a stable spatial separation between the sampling sites throughout the year, while seasonal differences were observed only within site-groups. This could be explained by the salinity gradient along the transect, as indicated by the high correlation with salinity (Fortunato et al. 2011). However, in absence of particular salinity gradients, also Wang et al. (Wang et al. 2019) found a strong influence of sample location on microbial community structure when sampling off the coastal PICO station and out to the Sargasso Sea. Yet, a seasonal effect could be seen there in the clustering of shelf stations, which grouped either with the coastal station or with the offshore station, depending on season and by the strong influence of temperature. Key differences between the study by Wang et al. (Wang et al. 2019) and our study are the strength of the environmental gradient between the coastal and offshore stations and the overall productivity of the coastal area. The region studied by Wang et al. (Wang et al. 2019) was characterized by higher productivity and stronger shore influence on the coastal stations. In the Wang et al. study primary production at the coastal station was usually above 1000 mg C m^−3^ d^−1^, which is more than an order of magnitude higher than that of their most offshore station (Wang et al. 2019). In the EMS, at a coastal site in Israel, Raveh et al. found that primary production would reach a maximum of about 270 mg C m^−3^ d^−1^ and the difference between coastal and offshore sites was only about four-fold (Raveh et al. 2015). Given that this site was only 50 m from shore with a bottom depth of 5m, we expect the difference in primary production between our coastal site (16 km from shore with a bottom depth of 100 m) and the offshore stations to be even lower. The spatial effects on the microbial community between the most coastal station 1 and the offshore stations 2 to 4 were detected in early winter and summer, yet not in spring, when coastal and offshore communities were highly similar. This is in line with earlier observations of the EMS, that based on chlorophyll and temperature, described a distinct border between coastal and offshore waters from late spring to early winter, while this border became diffuse from winter to early spring (Berman et al. 1986).

Profound and predictable seasonal shifts in marine surface water microbial communities have been shown in several studies (*e.g.* (Fuhrman et al. 2006, 2015; Ward et al. 2017; Galand et al. 2018)) and, as in our case, temperature has been frequently identified as a main environmental driver (*e.g.* (Sunagawa et al. 2015; Lucas et al. 2016; Ward et al. 2017)). Salinity, like temperature, is often an indicator for seasonal changes in hydrography and accordingly it is often identified as seasonal driver of marine surface water microbial communities (*e.g.* (Fuhrman et al. 2006; Ward et al. 2017)). Despite the fact that seasonality is expected to strongly affect primary productivity and nutrient availability in the EMS (Kress and Herut 2001; Krom et al. 2014; Raveh et al. 2015), our measured nutrient data explained the seasonal shifts in microbial communities to a lesser extent than the measured physical parameters. It is possible that the effect of nutrients was not detected as several measurements, especially phosphate, were below the detection limit of our analysis method. Higher sensitivity nutrient measurements using fresh unfrozen samples (such as those here utilized only in the July 2016 cruise) might clarify the importance of nutrients *versus* physical factors in the seasonality patterns of the microbial community. Our measured parameters explained a large part of the observed variability in microbial community: 49% of the DNA and 72% of the RNA data. Additional factors that were not measured in this study, may be responsible for the unexplained variability. These include UV radiation, which is known to penetrate deep into the clear waters of the EMS with UV doses among the highest of all oceans (Tedetti and Sempéré 2006; Smyth 2011), and irradiance, a factor known for structuring surface water microbial communities in the subtropical North Pacific Gyre station ALOHA (Bryant et al. 2016).

The recurrent shifts in microbial structure and activity followed established seasonal patterns. Physical mixing of the water column in early winter seemed to be responsible for a ‘resetting’ of the microbial ecosystem from a low diversity state in summer, when the microbial community is nutrient depleted in the EMS community (Thingstad et al. 2005; Tanaka et al. 2011), to a high diversity state in winter, as previously proposed for the Northwestern Mediterranean Sea (Salter et al. 2015). The mixing leads to an upwelling of nutrients that allow previously rare microbes to grow and thrive. In addition, microbes from deeper layers are brought up and might repopulate and grow in the now nutrient enriched surface waters as observed in the Western Mediterranean Sea (Haro-Moreno et al. 2018). These ‘upwelled’ microbes might interact with surface water microbes and affect the microbial community as shown in recent mesocosm experiments performed in the EMS (Hazan et al. 2018). Potential candidates of ‘upwelled’ microbes in our study include *e.g.* SAR202 and SAR406 groups that are typical of deeper water and were more abundant in early winter than in spring or summer. In the EMS, phytoplankton is dominated by nano- to micro sized organisms in winter and picophytoplankton in summer (Raveh et al. 2015). These differences were reflected in our pigment data. Seasonality in heterotrophic bacteria might thus be due to changes in phytoplankton composition, which, likely through the different types of organic carbon produced, are known to affect microbial community structure (Camarena-Gómez et al. 2018).

Annual microbial patterns can be observed also at the level of closely related OTUs within specific taxonomic groups that have different seasonal preferences, as found both in this study and at the coastal PICO site (Ward et al. 2017). The seasonal preference is likely due to differences in their genomic make-up resulting in different ecotypes that thrive under different environmental conditions. An example is provided by the dominant SAR11 clade, where, within clade 1a, the three most abundant OTUs peaked in summer, while the fourth to sixth most abundant ones peaked in spring. Nevertheless, the main seasonal differences in SAR11 abundances were linked to SAR11 clades. They followed the general patterns found at other ocean stations such as the Mola station in the Northwestern Mediterranean (Salter et al. 2015) and the BATS station in the Sargasso sea (Carlson et al. 2009; Vergin et al. 2013) with some differences: similarly to Mola and in contrast to BATS, in the EMS clade Ib did not replace clade Ia as the dominant clade in spring, whereas in contrast to Mola we found in the EMS a substantial amount of SAR11 clade IV that followed the seasonal pattern described at BATS (Vergin et al. 2013).

Seasonal shifts in microbial communities are linked to functional differences between microbes that enable better adaptation to the changing environment (Galand et al. 2018; Haro-Moreno et al. 2018). These changes can have a strong impact on biogeochemical cycles. Here we provide a detailed analysis of spatial and temporal changes of microbial communities in the South Eastern Levatine basin. To understand how the observed microbial community shifts affect biogeochemial cycles will require metagenomic and transcriptomic studies, as well as ecophysiology investigations of the dominant bacterial groups.

## Acknowledgements

We thank all cruise participants, the crew of the R/V Mediterranean Explorer and the EcoOcean foundation for their help in sampling, Dr. Rinat Bar Shalom (University of Haifa) for help in cruise planning and preparation, Dr. Tanya Rivilin (Interuniversity Institute for Marine Science at Eilat) for help in nutrient analysis and Dr. Stephan Green (DNA Services Facility at University of Illinois at Chicago) for useful comments and suggestions on the amplicon sequencing methods. This study was funded by the Israel Science Foundation grant (ISF #1243/16) to LS. The seasonal cruises were supported by funding from the Leon H. Charney School of Marine Sciences (Haifa University, Israel). MH was supported by an Inter-Institutional post-doctoral fellowship from the Haifa University and a Helmsley Trust fellowship.

