## Supplementary Figures and Tables for "Microbial communities in an ultra-oligotrophic sea are more affected by season than by distance from shore"

For the article:

Authors’ affiliations:

^1^ Marine Microbiology Lab, Department of Marine Biology, Leon H. Charney School of Marine Sciences, Haifa University

^2^ Department of Aquatic Microbial Ecology, Institute of Hydrobiology, Biology Centre CAS, Czech Republic

^3^ Bioinformatics Service Unit, University of Haifa

^4^ The Dr. Moses Strauss Department of Marine Geosciences, Leon H. Charney School of Marine Sciences, Haifa University

^5^ Morris Kahn Marine Research Station, Environmental Geochemistry Lab., Leon H. Charney School of Marine Sciences, Haifa University

Supplementary Figure S1 A: Temperature - depth (up to 500 m) profiles of the transect based on CTD data. All CTD plots were obtained with odv4 (Version 4.7.4) (Schlitzer, R. 2015. Ocean Data View. http://odv.awi.de). Data were stretched by weighted-average gridding with automatic scale length. Bad estimates were hidden, color shading enabled and a quality limit of 5.0 applied. Contours indicate increments of 1 °C. Stations are indicated by vertical lines, order from left to right: Station 1, 1A, 2, 2B, 3, and 4. Color scale is the same for all plots. X-axis indicates distance from shore.

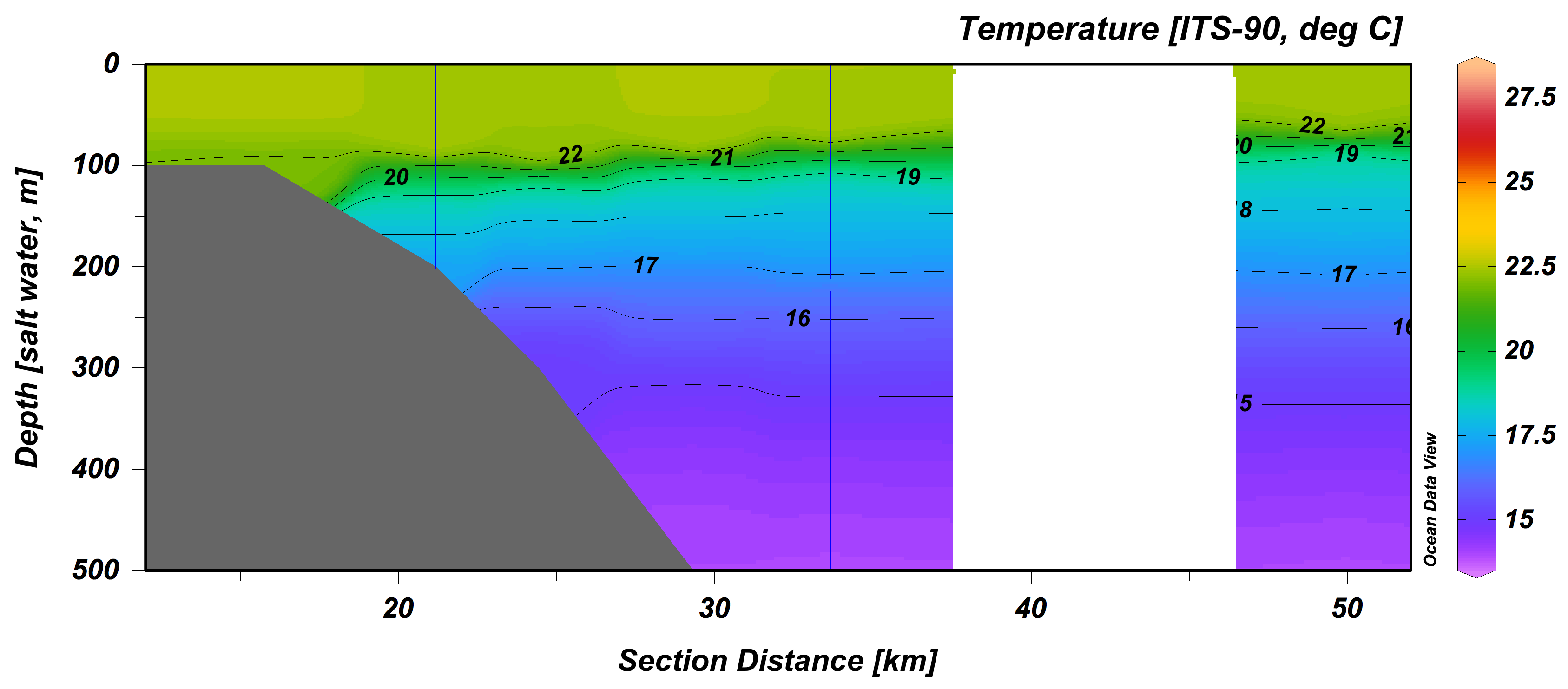

Spring 2015

Winter 2014

Summer 2015

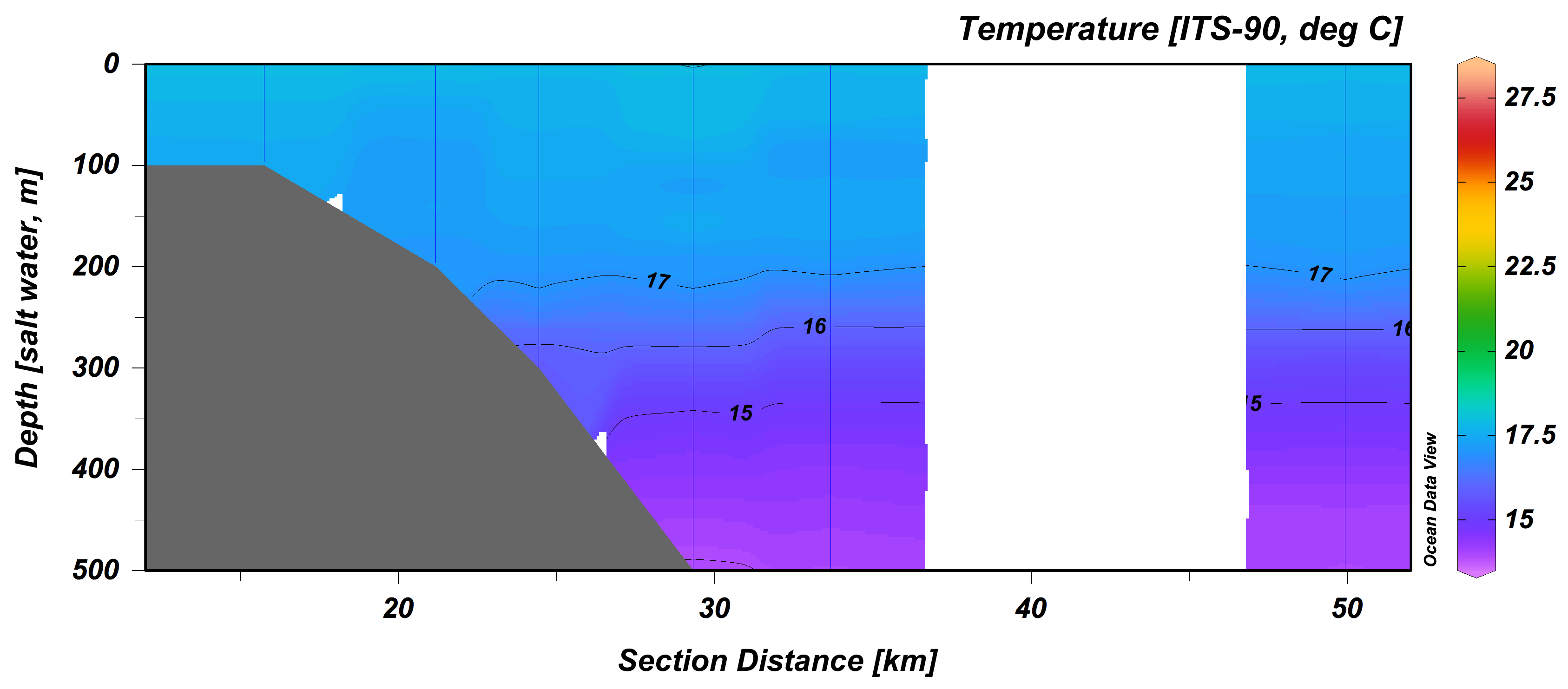

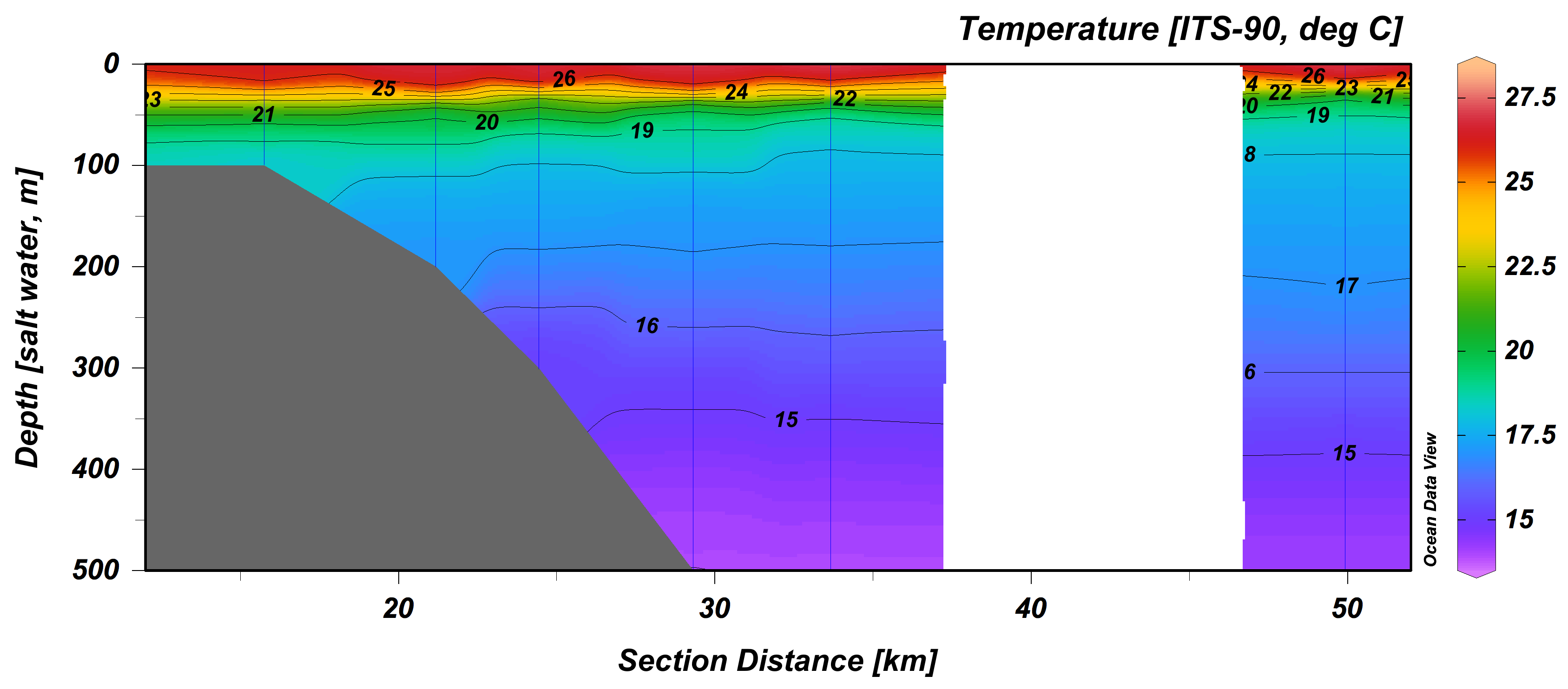

Supplementary Figure S1 A: continued.

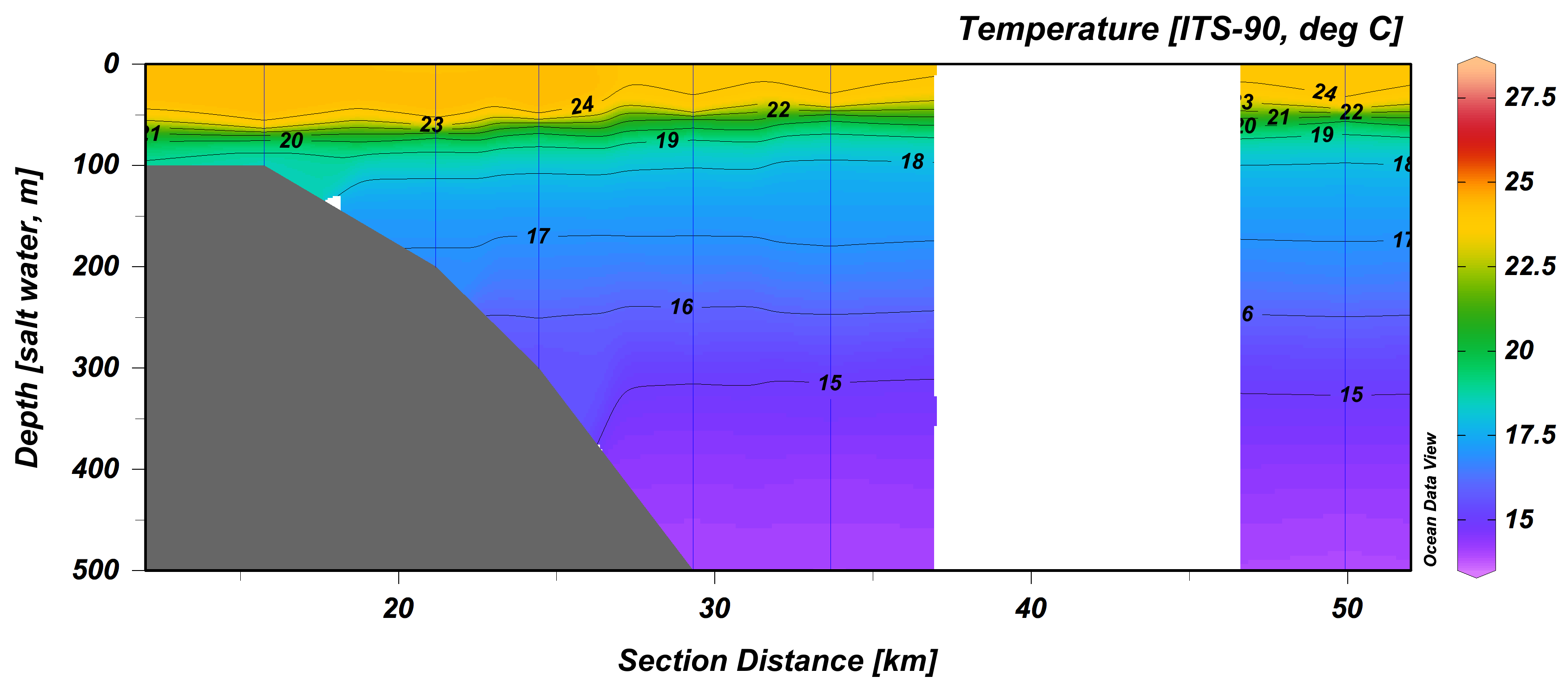

Spring 2016

Winter 2015

Summer 2016

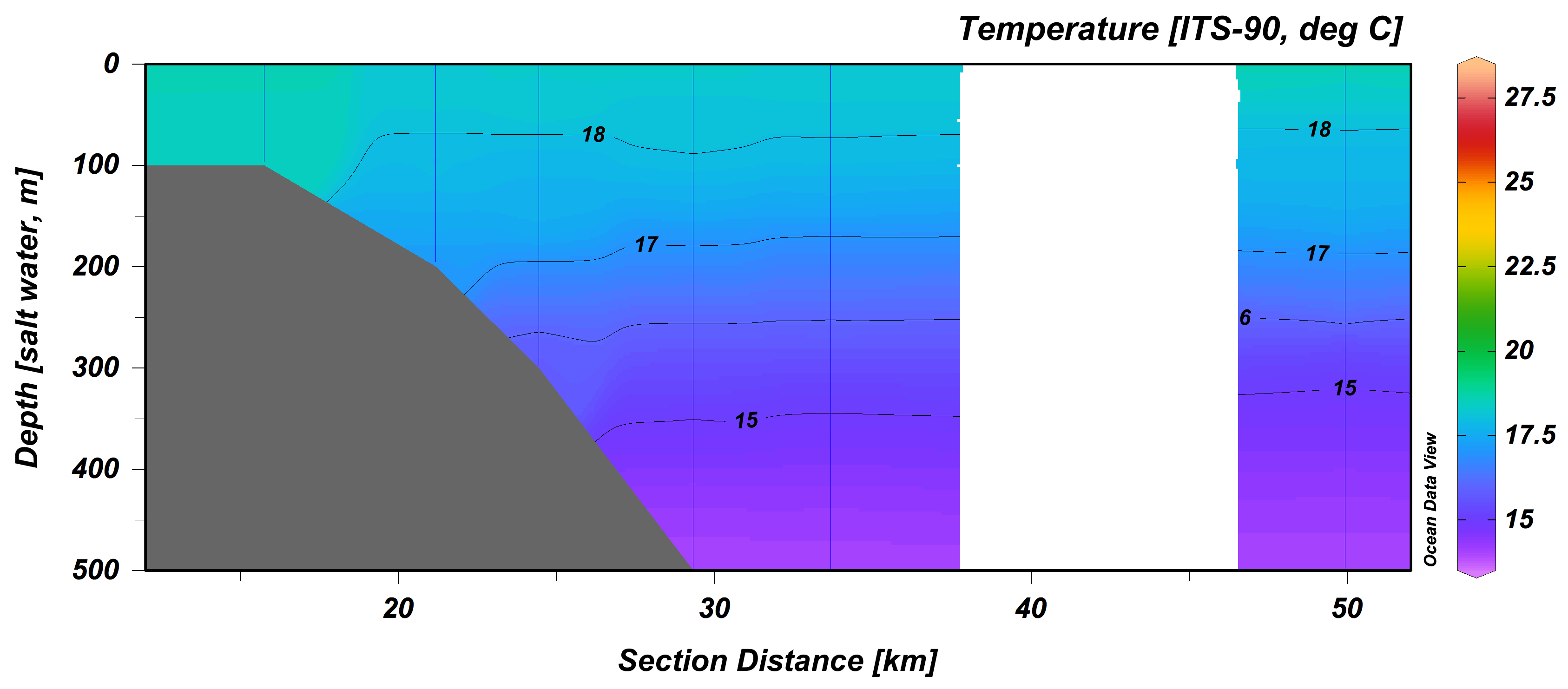

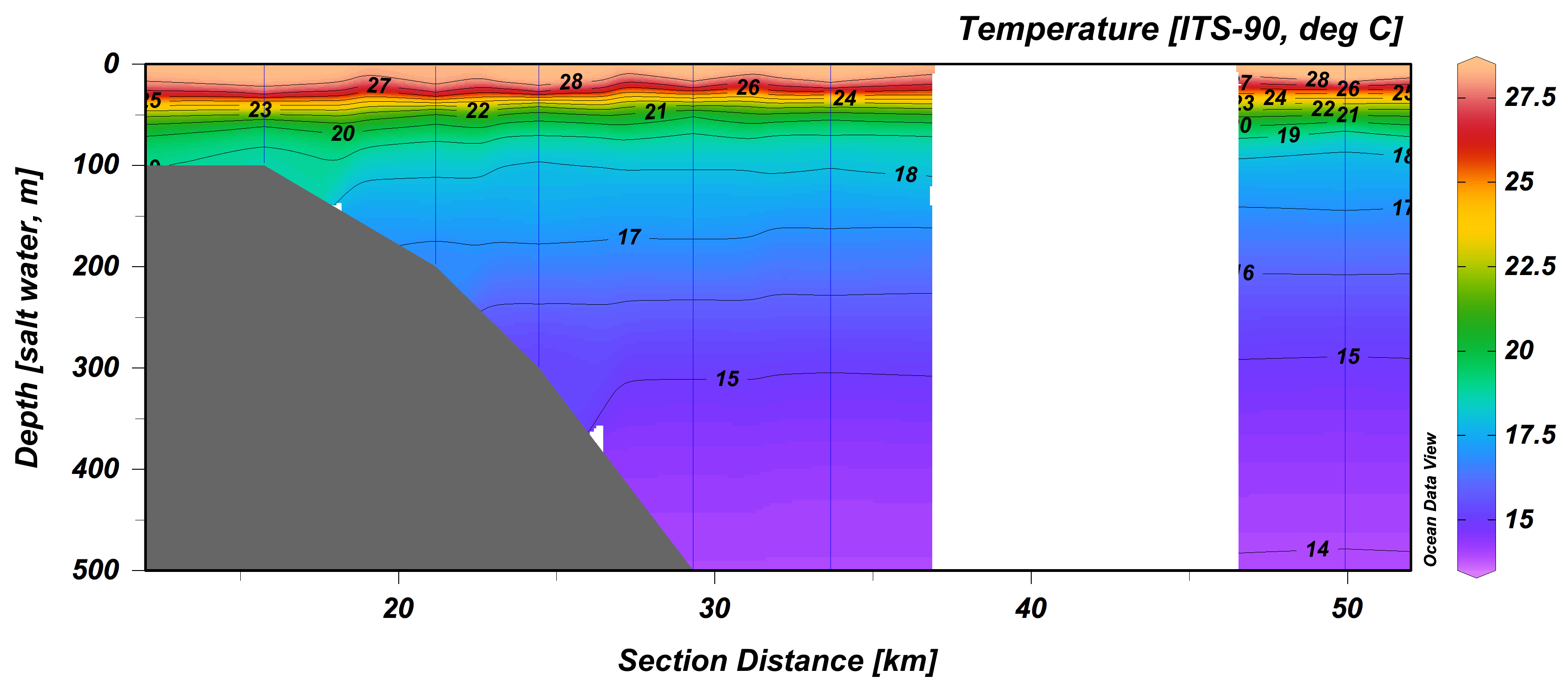

Supplementary Figure S1 B: Salinity - depth profiles of based on CTD data. Due to technical problems, no salinity data were available for station 1 in the spring 2016 cruise. For software settings see A. Contours indicate increments of 0.05 PSU. Color scale is the same for all plots.

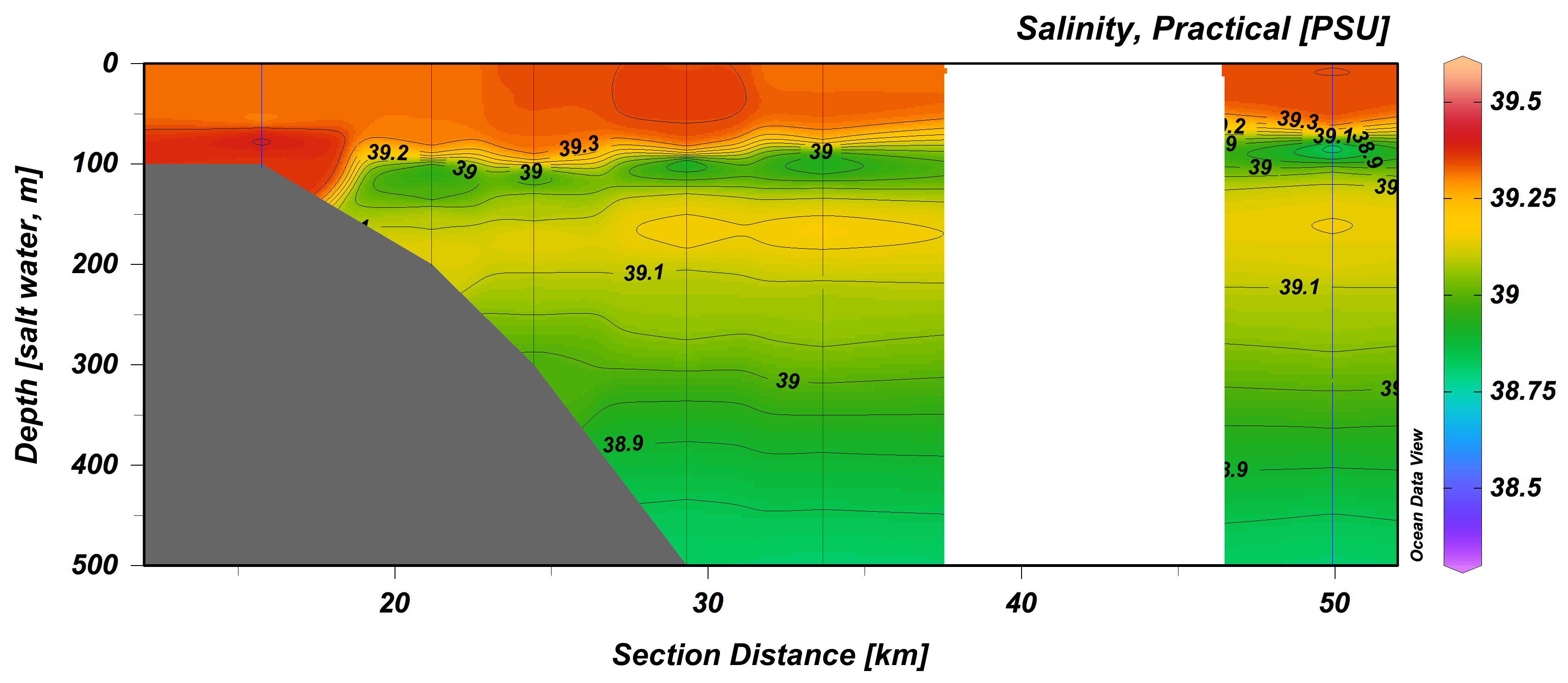

Spring 2015

Winter 2014

Summer 2015

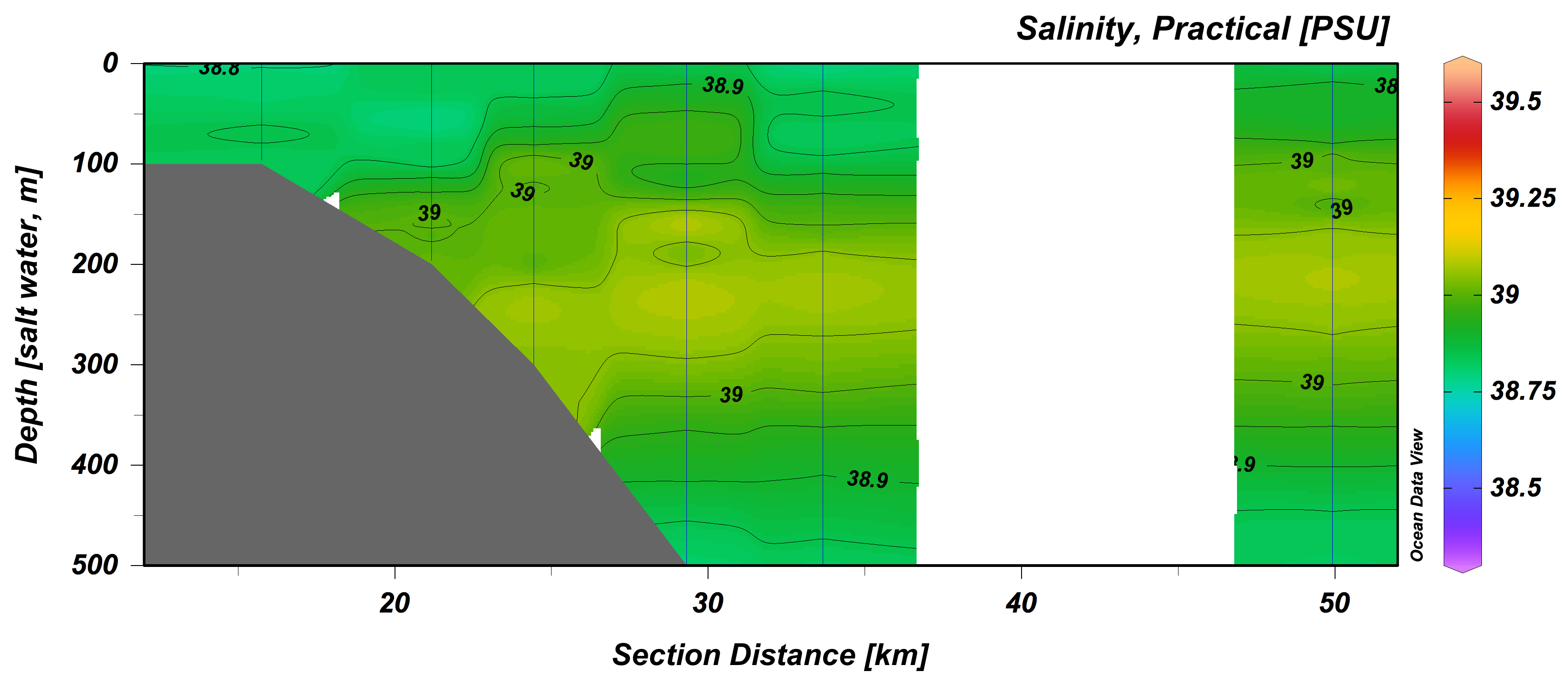

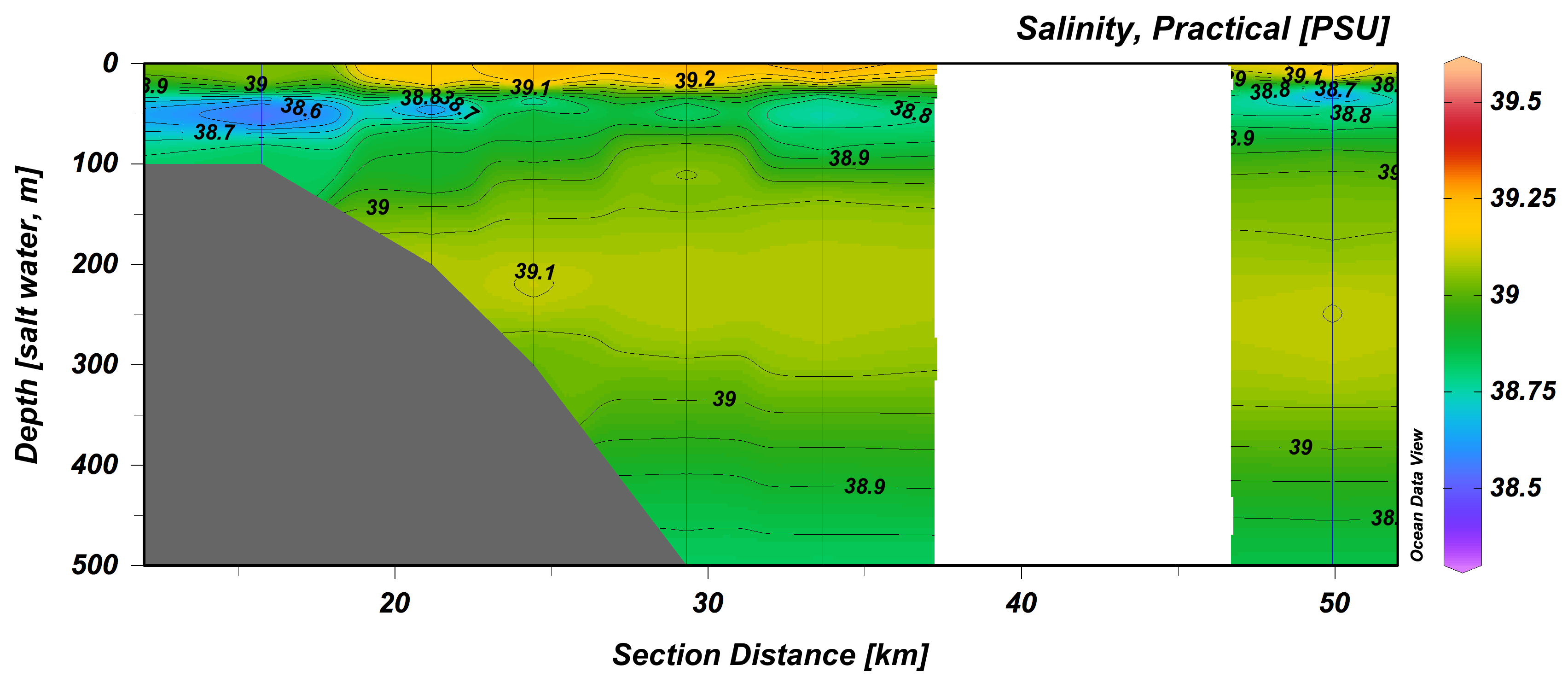

Supplementary Figure S1 B: continued.

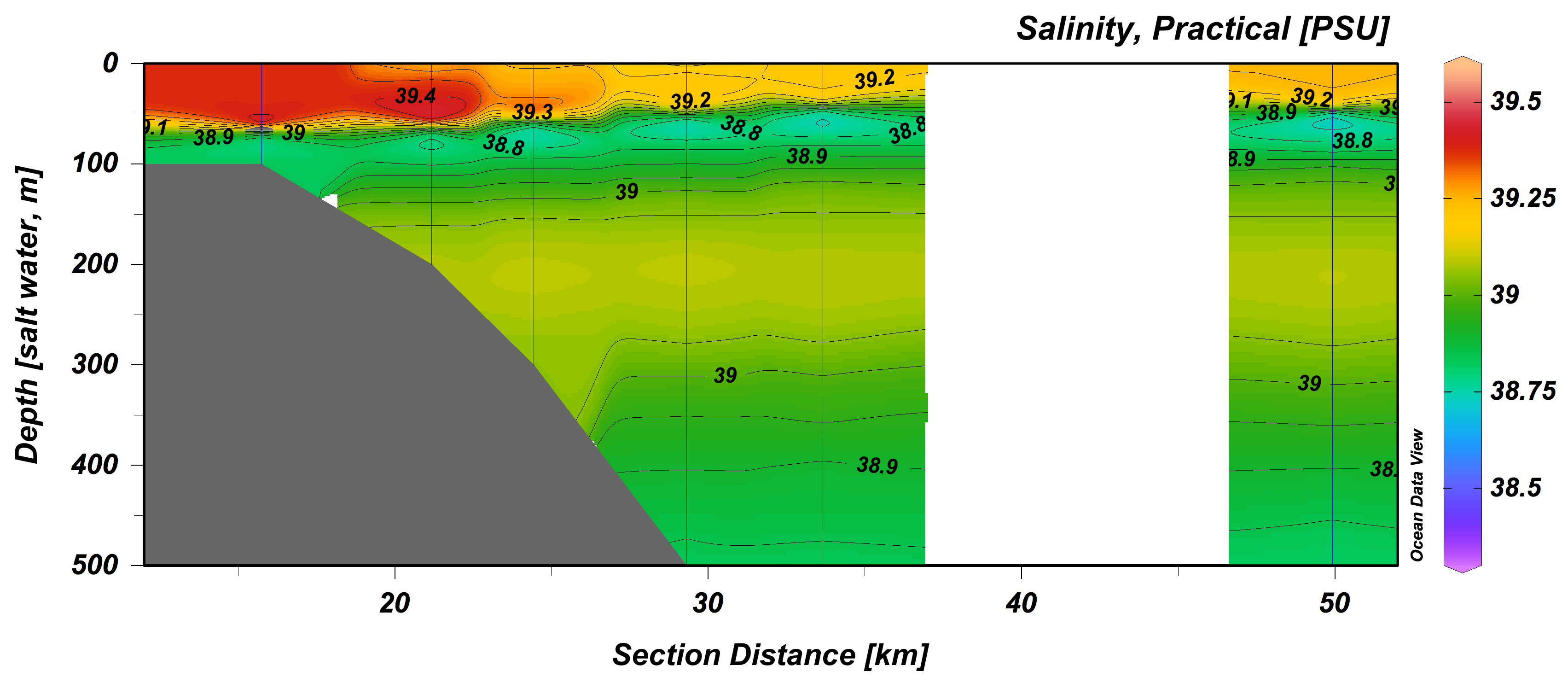

Spring 2016

Winter 2015

Summer 2016

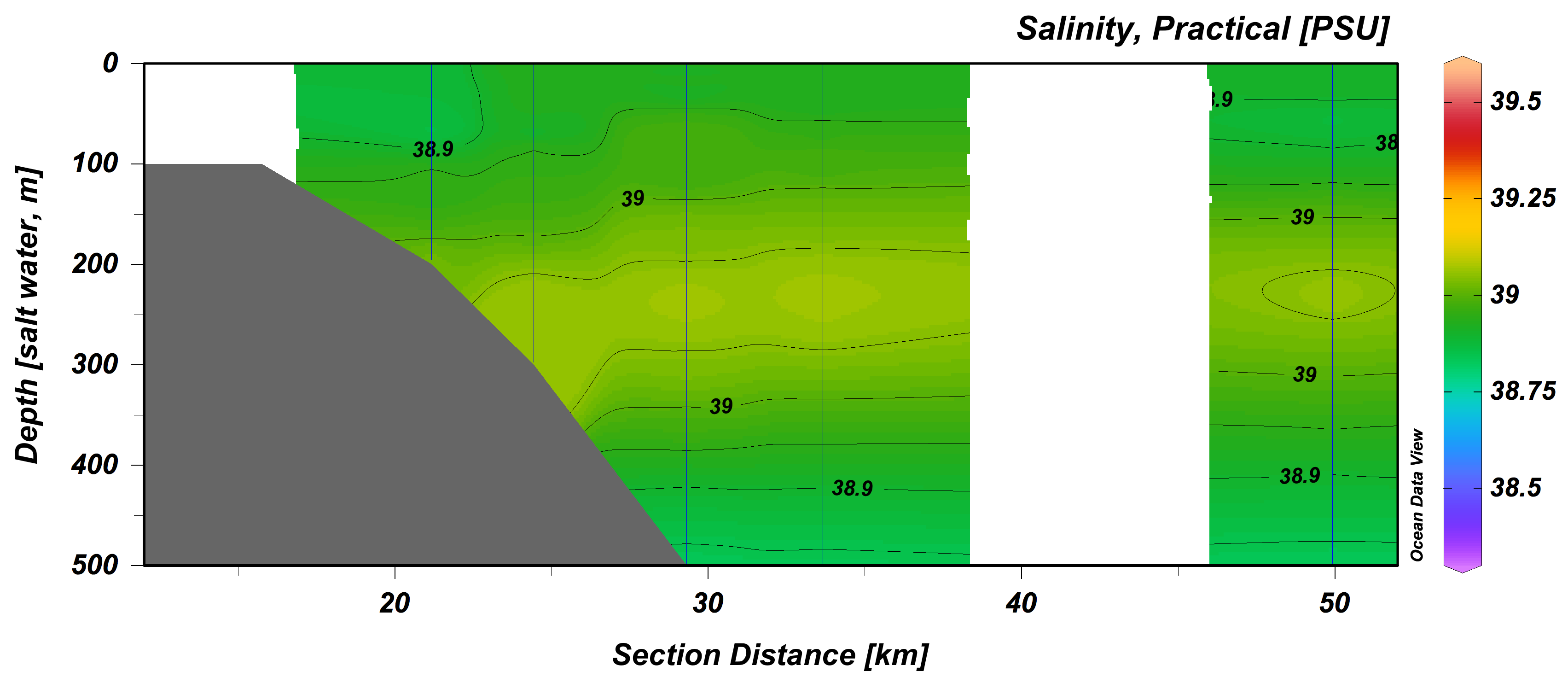

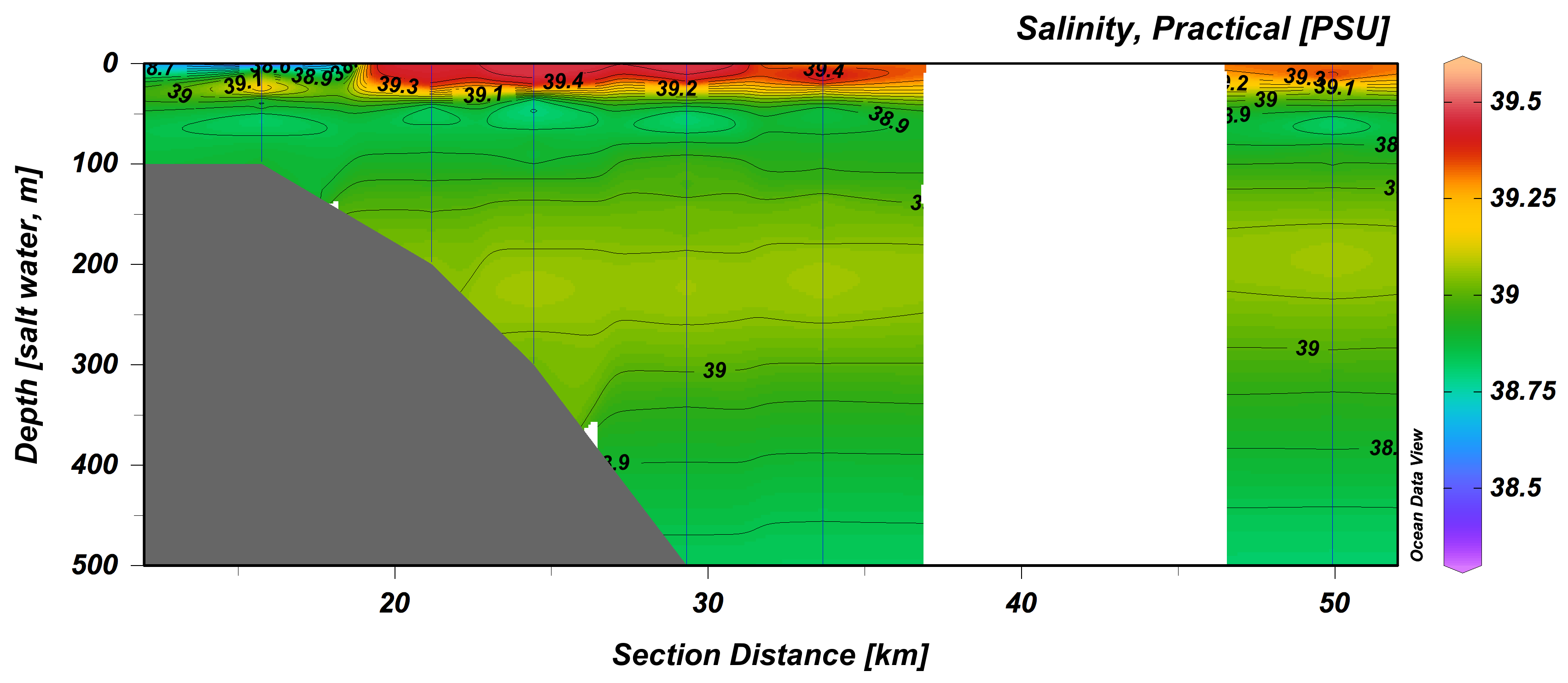

Supplementary Figure S1 C: Chlorophyll *a* fluorescence – depth profiles based on CTD data. For software settings see A. Contours indicate increments of 0.05. Color scale is the same for all plots.

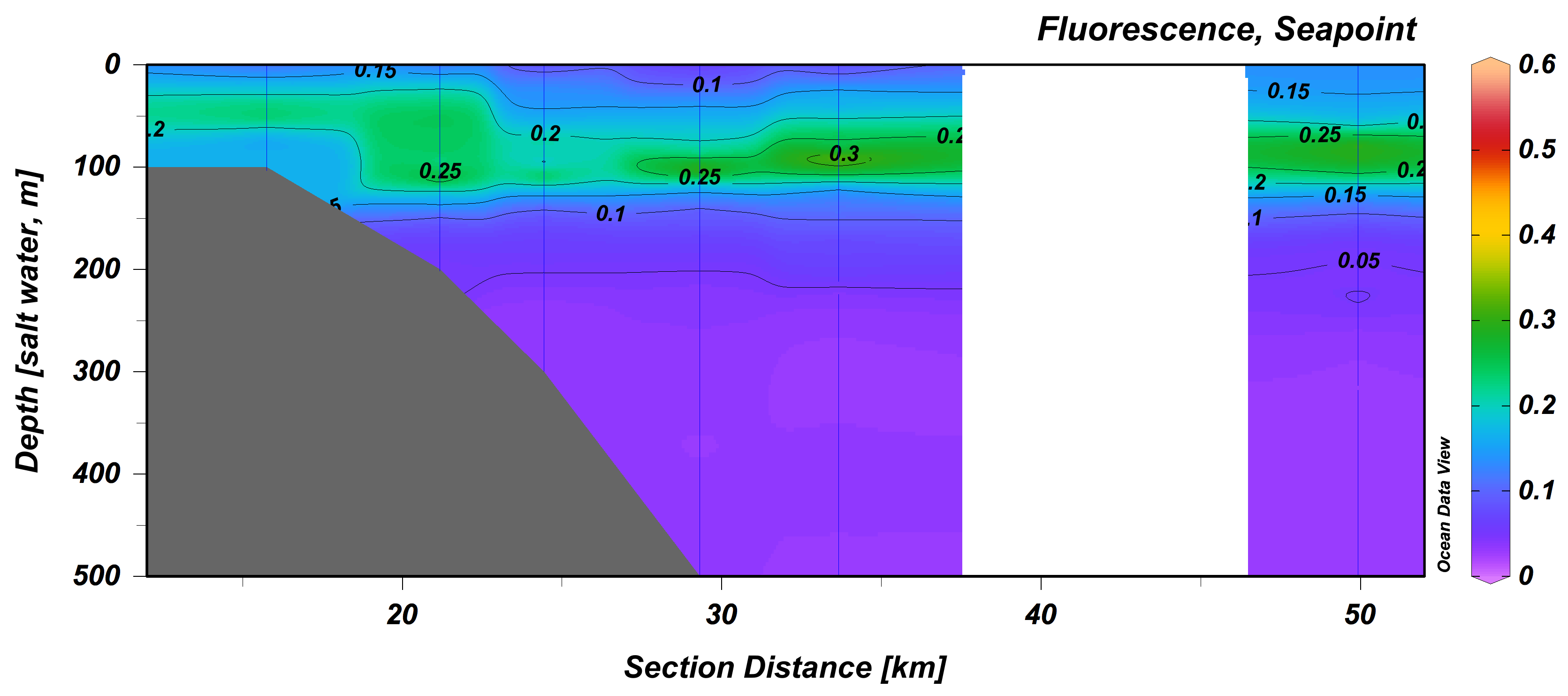

Spring 2015

Winter 2014

Summer 2015

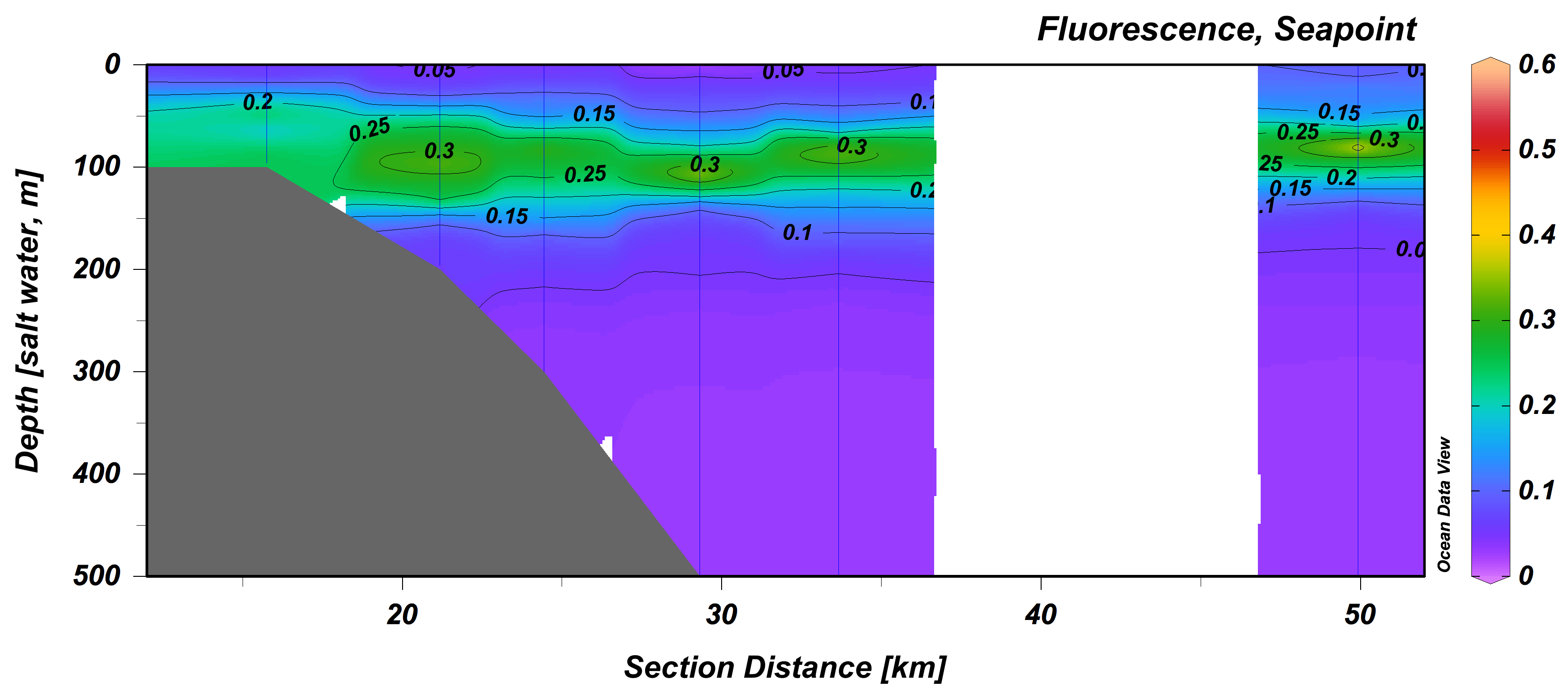

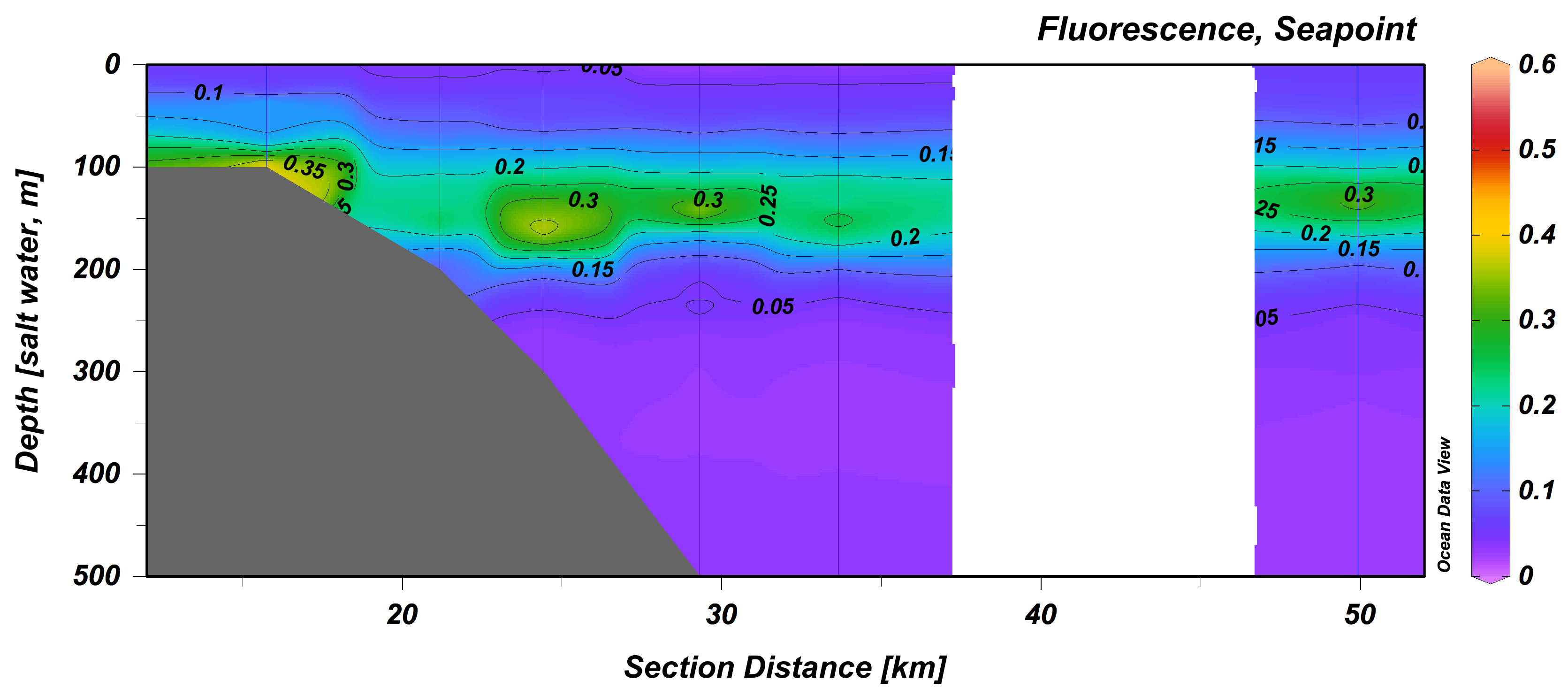

Supplementary Figure S1 C: continued.

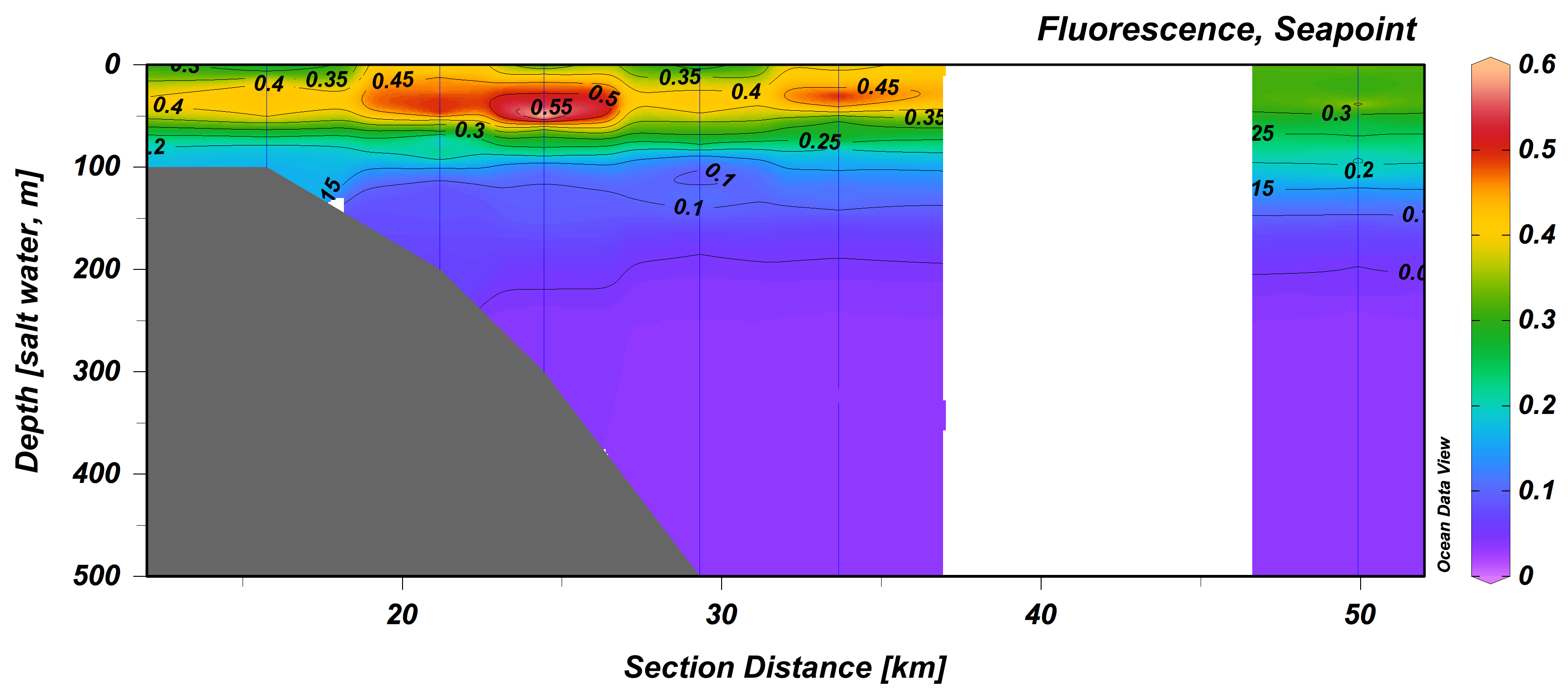

Spring 2016

Winter 2015

Summer 2016

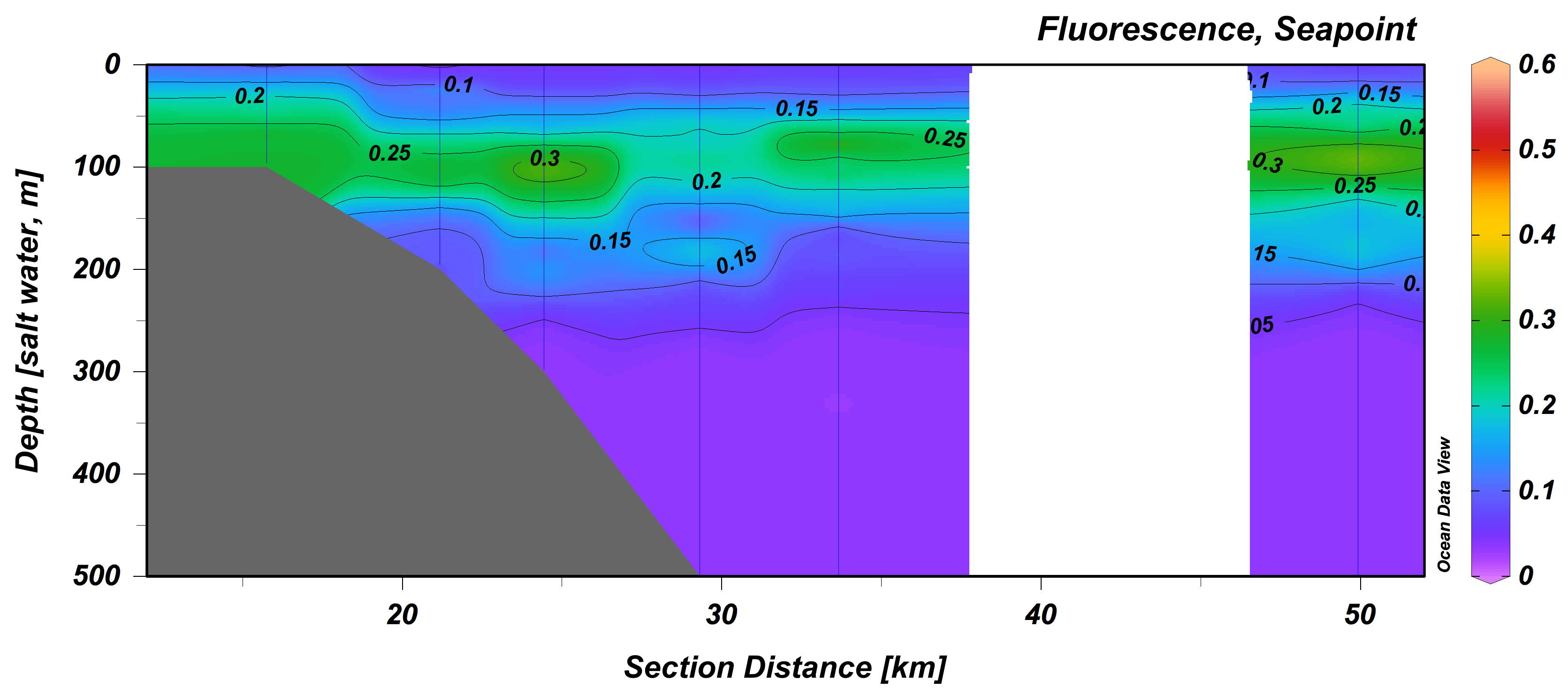

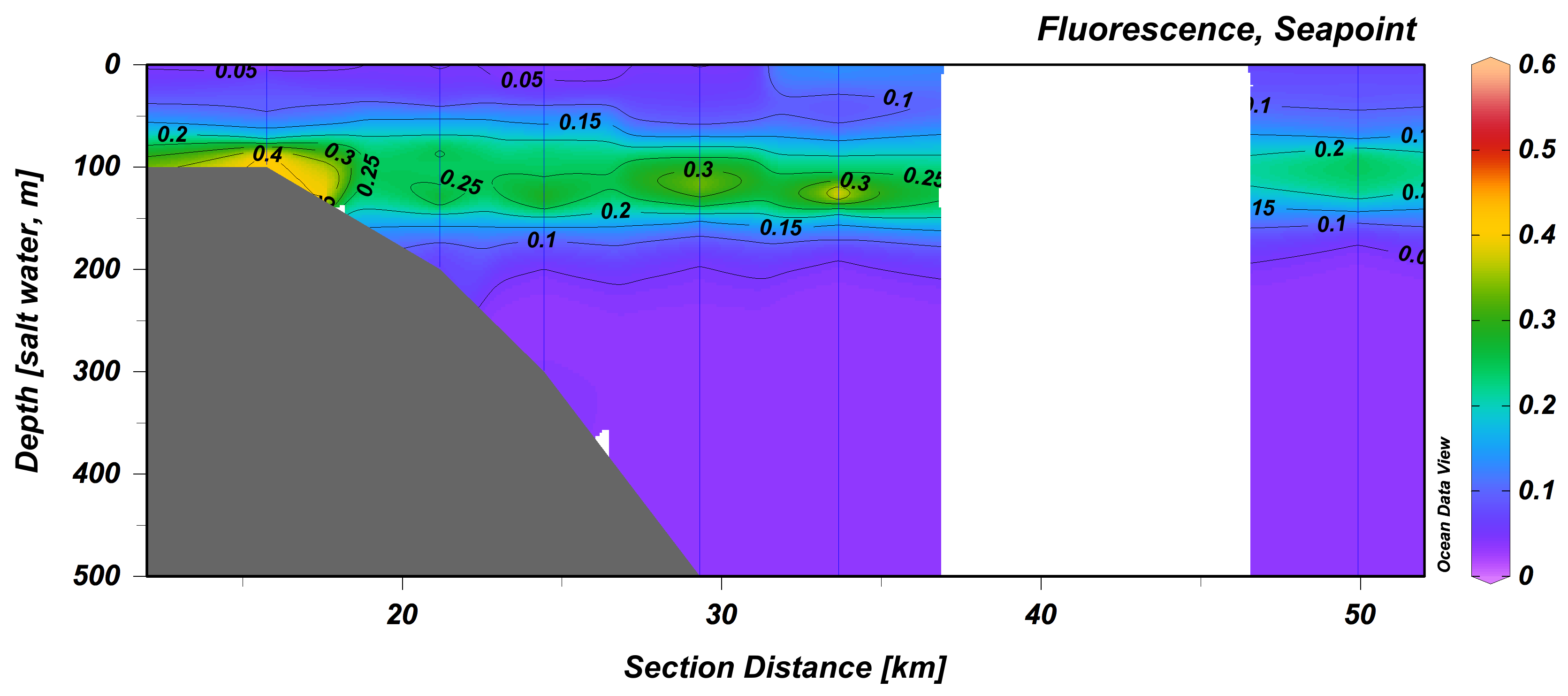

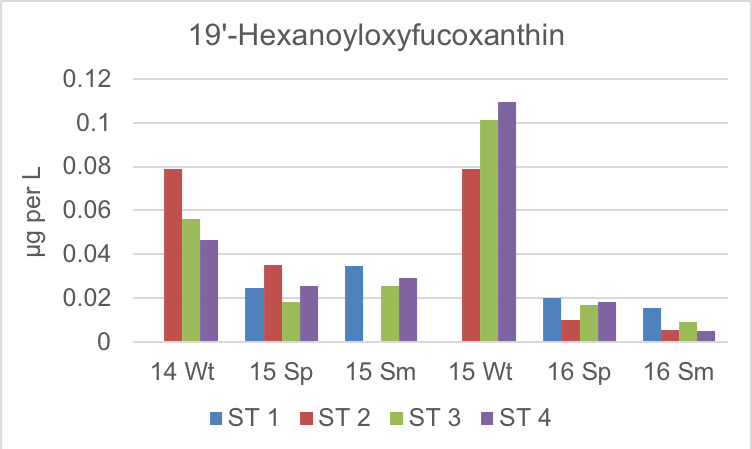

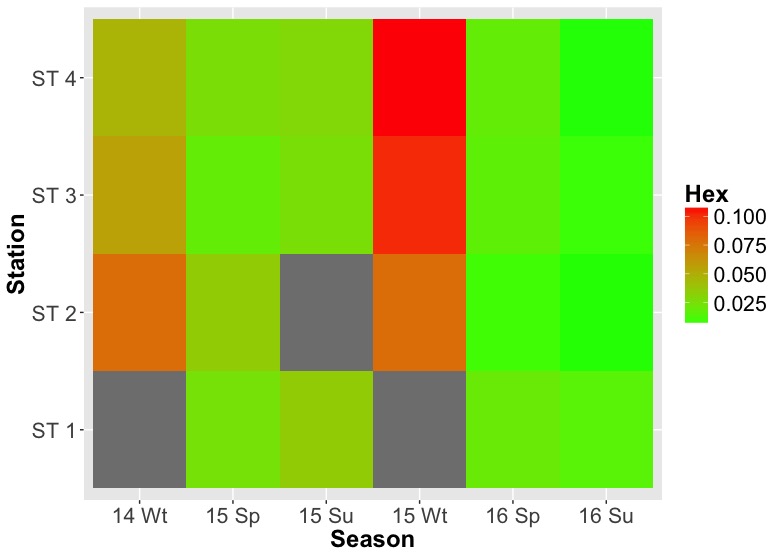

B

A

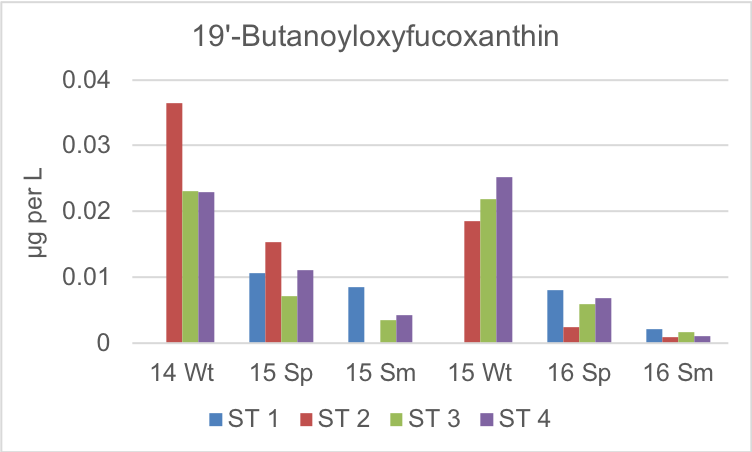

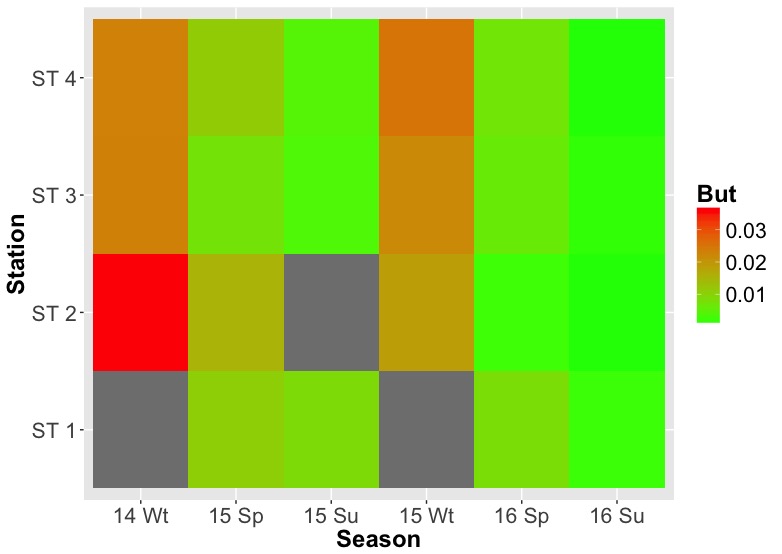

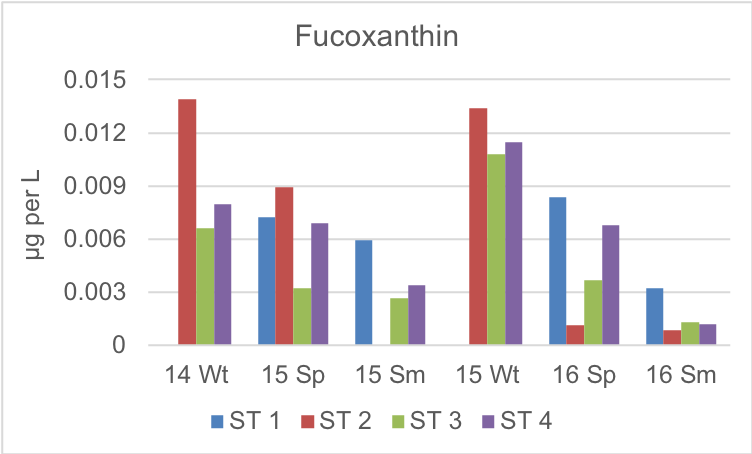

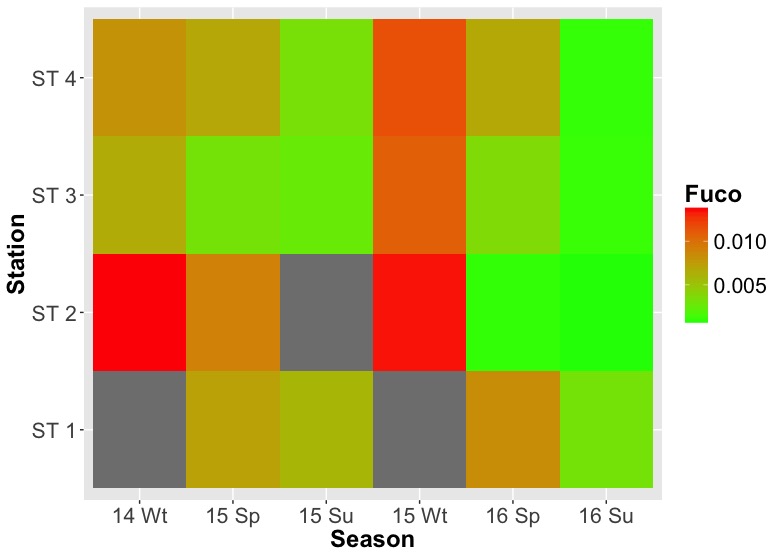

C

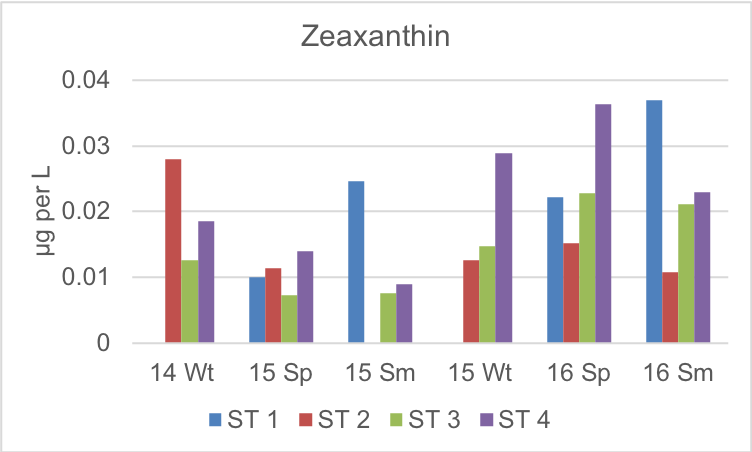

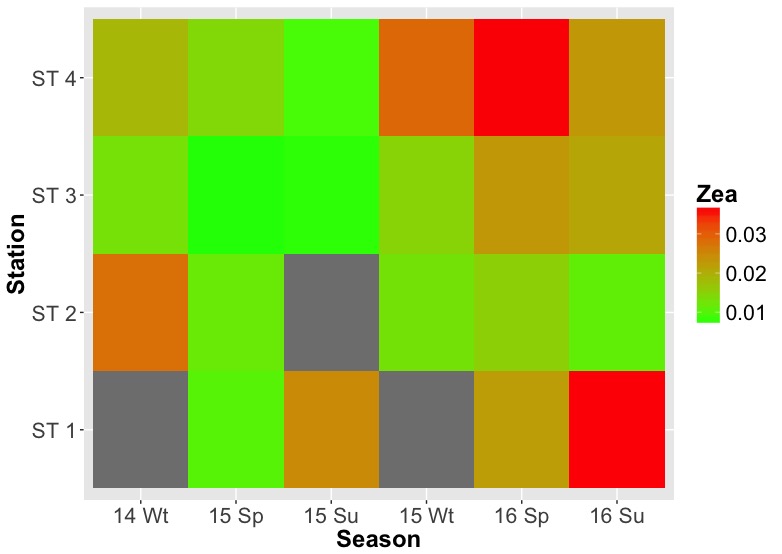

D

Supplementary Figure S2: Concentrations of the main detected diagnostic pigments: A) 19’-hexanoyloxyfucoxanthin, B) 19’-butanoyloxyfucoxanthin, C) fucoxanthin, D) zeaxanthin. No pigment samples were available for station 1 in early the winter 2014 and 2015 and for station 2 in the summer 2015 as indicated by the grey tiles in the heat maps.

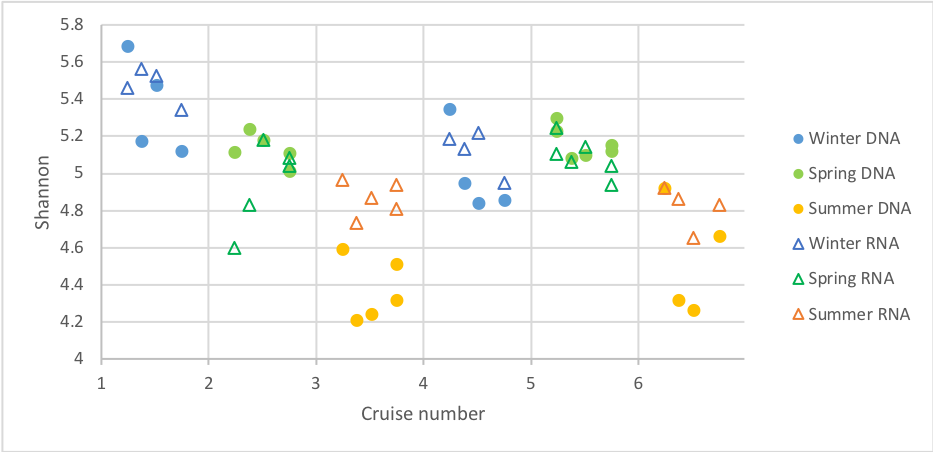

A

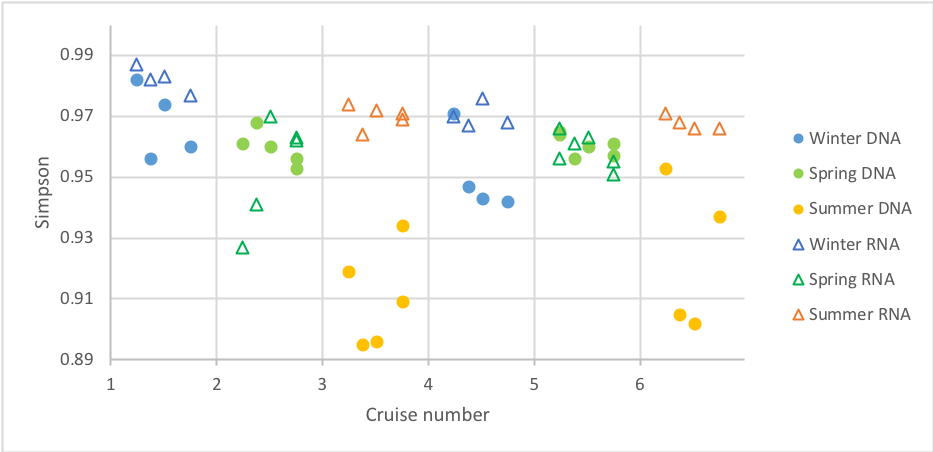

B

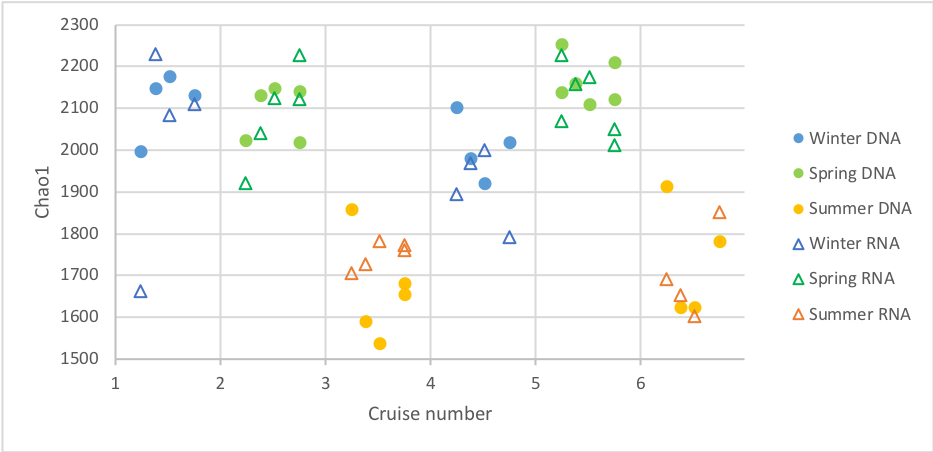

C

Supplementary Figure S3: Seasonal difference in diversity indices. A) Shannon, B) Simpson, C) Chao1. Stations are sorted from station 1 to 4 (left to right) along the x-axis within each cruise. Significant seasonal differences were found in all three alpha diversity indices for both resident and active communities (Kruskal-Wallis test, *P*<0.001 in all six seasonal comparisons). DNA samples: In all three indices summer samples were less diverse than spring and early winter samples (Dunn’s post hoc test, Bonferroni corrected *P*<0.05), with no difference between the latter two.

In RNA samples: Shannon diversity was significantly lower in summer samples than early winter samples (Dunn’s post hoc test, Bonferroni corrected *P*<0.05) with spring samples not differing from summer nor early winter samples; Chao1 was lower in summer samples than in spring samples (Dunn’s post hoc test, Bonferroni corrected *P*<0.05) with early winter samples not differing from summer nor spring samples; Simpson diversity was lower in spring samples compared to summer and early winter samples (Dunn’s post hoc test, Bonferroni corrected *P*<0.05) with no differences between the latter two.

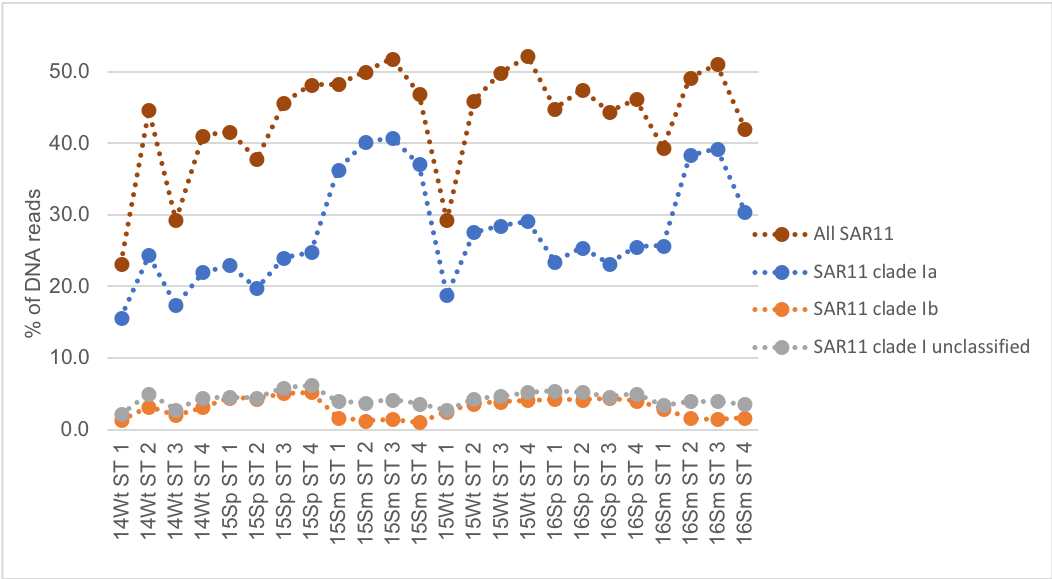

A

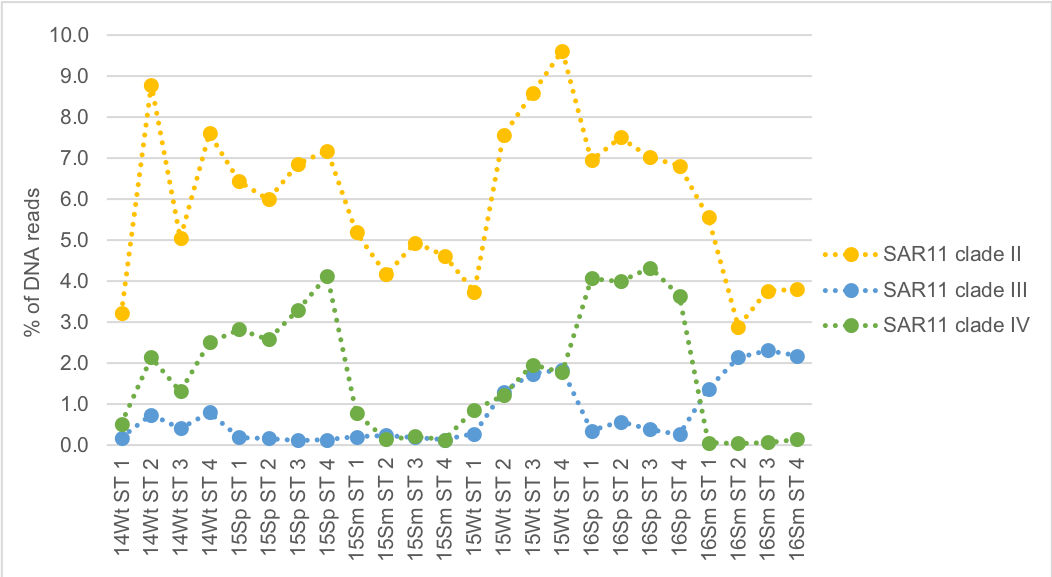

B

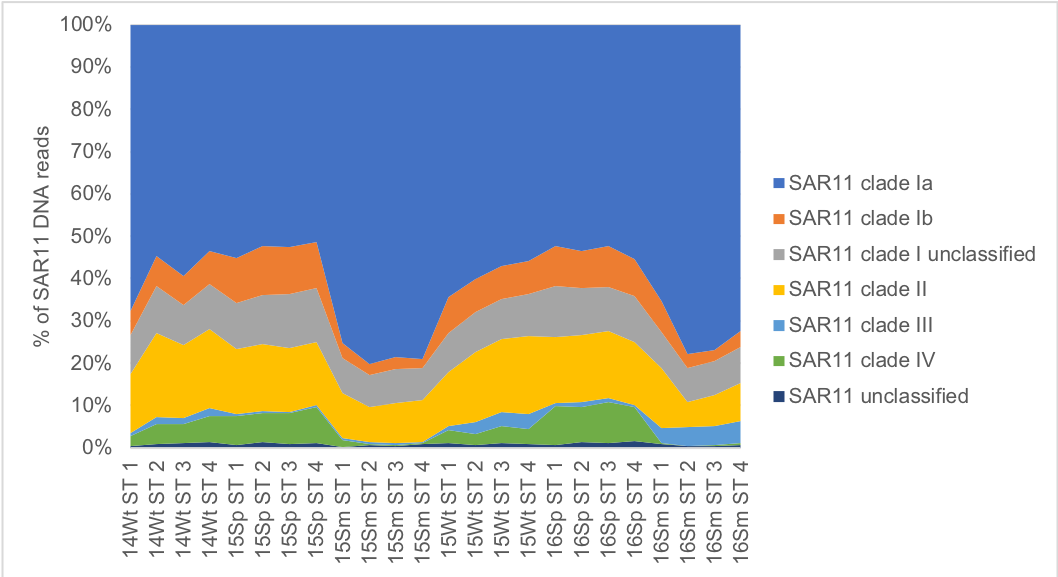

C

Supplementary Figure S4: Seasonality of SAR11 clades according to DNA. Relative abundances of: A) SAR11 order and clade 1 subclades; B) non-clade 1 SAR11 clades. C) Contribution of different clades to all SAR11 DNA reads.

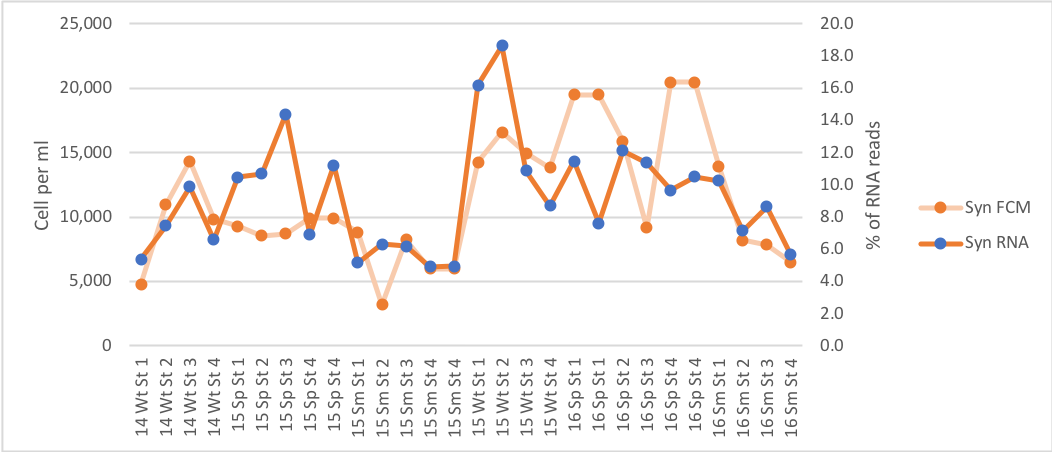

A

B

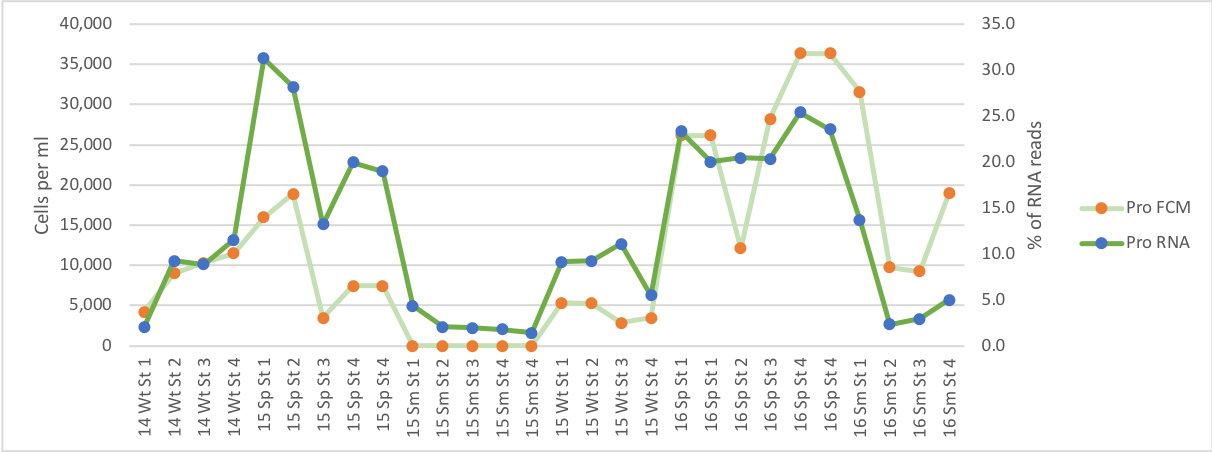

Supplementary Figure S5: Seasonal activity and abundance of A) *Synechococcus* and B) *Prochlorococccus* based on relative abundance of RNA reads and cell counts from flow cytometry (FCM).

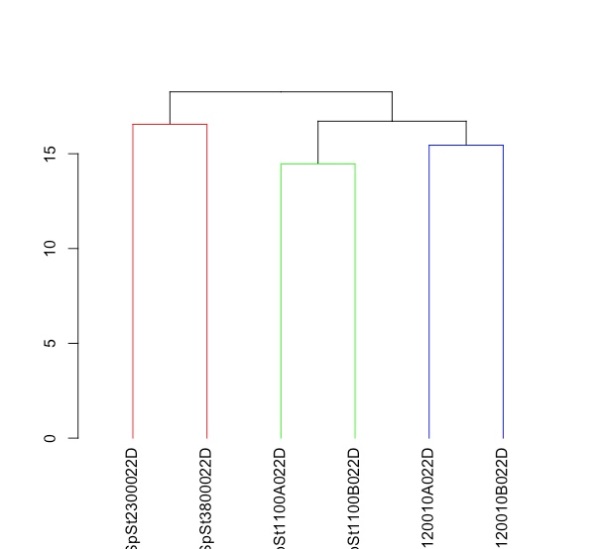

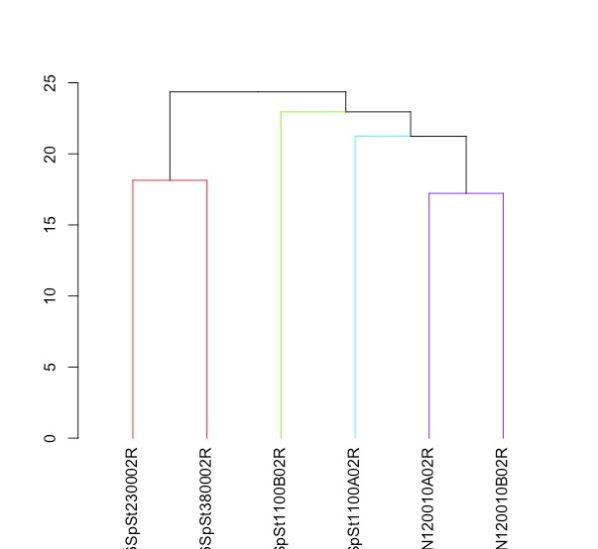

Bray-Curtis dissimilarity %

ST 2

ST 3

ST 1 R2

ST 1 R1

ST 4 R1

ST 4 R2

ST 2

ST 3

ST 1 R1

ST 1 R2

ST 4 R1

ST 4 R2

Bray-Curtis dissimilarity %

B

A

Supplementary Figure 6: SIMPROF clustering results for A) DNA and B) RNA samples from the spring 2016 cruise. R1 and R2 indicate replicates from the same location. Distances were based on Bray-Curtis dissimilarities and clusters were linked based on average-linkage. Significant clusters (alpha=0.05) are colored.

Direct analyses of the Bray-Curtis dissimilarities confirmed small, significant differences between the stations affected by the intrusion (stations 1 and 4) and not affected (stations 2 and 3) (in DNA samples average dissimilarity between replicates of stations 1 and 4: 0.167, between station 2 and 3: 0.166 and average between station 1 & 4 and station 2 & 3: 0.183; RNA: 0.216, 0.181, 0.244) (Mann-Whitney U test, z=-2.4627, *P*=0.014 for DNA and z=-2.123, *P*=0.034 for RNA).

DNA (variance explained 49%)	RNA (variance explained 72%)

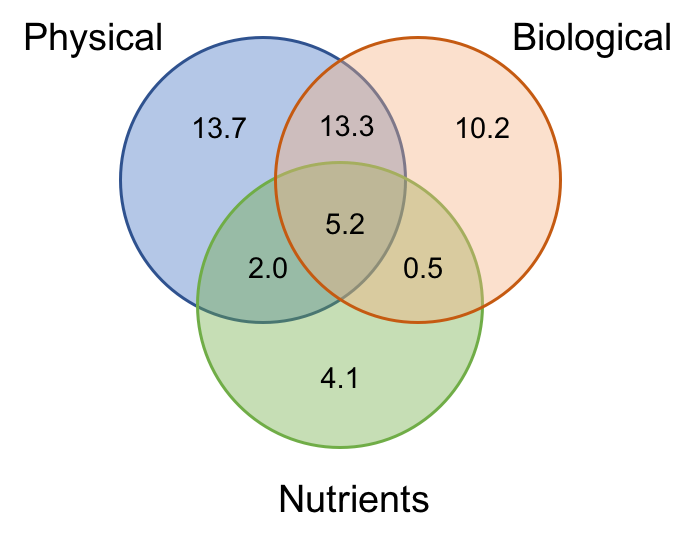

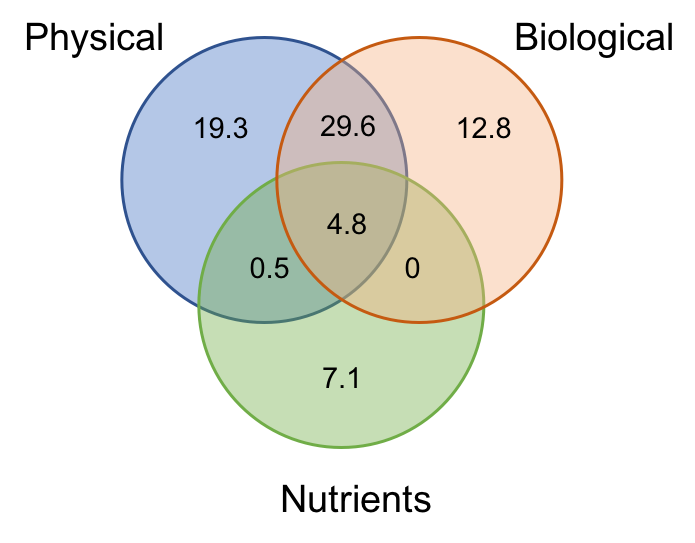

B

A

| Analysis |  | DNA | | | RNA | | |
| --- | --- | --- | --- | --- | --- | --- | --- |
|  |  | Physical | Biological | Nutrients | Physical | Biological | Nutrients |
| Variation partitioning | Total VE (%) | 34.2*** | 29.2*** | 11.8*** | 54.2*** | 45.1*** | 10.3** |
|  | Independent VE (%) | 13.7*** | 10.2* | 4.1*** | 19.3** | 12.8 * | 7.1 ns |
| CCA | Inertia explained (%) | 33.3*** | 32.5*** | 25.2*** | 36.4*** | 30.3*** | 21.2** |
|  | Significant variables (P<0.05) | Temperature; Salinity | Fluorescence; Total cell count;  *Prochlorococcus* cell count | Phosphate;  NO3+NO2;  Silicate | Distance between stations; Temperature; Salinity;  Turbidity | Fluorescence; Total count;  *Prochlorococcus* cell count | Silicate |
| Partial CCA (conditioned) | Inertia explained (%) | 15.6*** | 15.4*** | 13.5*** | 17.9** | 13.5* | 11.1 ns |
|  | Significant variables (P<0.05) | Temperature; Salinity | Fluorescence | Silicate | Temperature; Salinity | Fluorescence | ns |

Supplementary Figure S7: Variation in microbial community composition explained by the physical, biological and nutrient matrixes. A) DNA and B) RNA Venn diagrams of the explained variation by matrix based on the variation partitioning (RDA) analysis. Table gives the overview of significance and the significant factors in the CCA analysis and the partial CCA analysis conditioned by the other two matrices. Asterisks indicate significance level for the model: * p<0.05; ** p<0.01; ***p<0.001; ns not significant.

Supplementary Table S1: Station locations, bottom depth, distance to shore and distances between stations.

Station 1A and 2B were sampled to identify gradients in the CTD data and from the second cruise onwards also in the nutrients.

| **Station** | **Location**  **(NMEA)** | **Bottom depth (m)** | **Distance (km) to** | | | | | |
| --- | --- | --- | --- | --- | --- | --- | --- | --- |
|  |  |  | **Herzliya Marina** | **ST 1** | **ST 1A** | **ST 2** | **ST 2B** | **ST 3** |
| ST 1 | 32 15.24 N  034 39.53 E | 100 | 16.13 | * |  |  |  |  |
| ST 1A | 32 17.07 N  034 36.89 E | 200 | 21.48 | 5.35 | * |  |  |  |
| ST 2 | 32 18.26 N  034 35.07 E | 300 | 25.08 | 8.95 | 3.60 | * |  |  |
| ST 2B | 32 20.09 N  034 32.66 E | 600 | 30.15 | 14.02 | 8.67 | 5.07 | * |  |
| ST 3 | 32 21.32 N 034 30.62 E | 800 | 34.06 | 17.93 | 12.58 | 8.98 | 3.91 | * |
| ST 4 | 32 26.99 N  034 22.51 E | 1200 | 50.54 | 34.41 | 29.06 | 25.46 | 20.39 | 16.48 |

Supplementary Table S2: Nutrient data in µmol/l. Precision limits are 0.05 µmol/l for silica and nitrate+nitrite and 0.04 µmol/l for phosphate. Nutrients were measured in duplicates. From Spring 2015 onwards nutrients were also measured at the intermediate stations 1A and 2B to improve detection of potential gradients.

| Year Season | Station | Phosphate | | Nitrate+Nitrite | | Silicate | |
| --- | --- | --- | --- | --- | --- | --- | --- |
|  |  | Average | SD | Average | SD | Average | SD |
| 2014 Winter | ST 1 | 0.07 | 0.01 | 0.69 | 0.37 | 2.10 | 0.05 |
|  | ST 2 | 0.01 | 0.01 | 0.34 | 0.14 | 0.47 | 0.15 |
|  | ST 3 | 0.03 | 0.03 | 0.32 | 0.19 | 0.50 | 0.27 |
|  | ST 4 | 0.06 | 0.00 | 0.18 | 0.07 | 0.63 | 0.06 |
| 2015 Spring | ST 1 | 0.05 | 0.00 | 0.19 | 0.02 | 1.02 | 0.06 |
|  | ST 1A | 0.04 | 0.00 | 0.26 | 0.04 | 0.84 | 0.00 |
|  | ST 2 | 0.07 | 0.01 | 0.34 | 0.06 | 0.52 | 0.01 |
|  | ST 2B | 0.04 | 0.00 | 0.48 | 0.03 | 0.61 | 0.08 |
|  | ST 3 | 0.04 | 0.00 | 0.64 | 0.03 | 0.40 | 0.01 |
|  | ST 4 | 0.03 | 0.00 | 0.10 | 0.00 | 0.66 | 0.01 |
| 2015 Summer | ST 1 | 0.05 | 0.01 | 0.12 | 0.04 | 0.54 | 0.00 |
|  | ST 1A | 0.04 | 0.01 | 0.04 | 0.00 | 0.51 | 0.00 |
|  | ST 2 | 0.04 | 0.01 | 0.05 | 0.00 | 0.52 | 0.00 |
|  | ST 2B | 0.04 | 0.00 | 0.02 | 0.00 | 0.54 | 0.03 |
|  | ST 3 | 0.03 | 0.00 | 0.04 | 0.00 | 0.58 | 0.05 |
|  | ST 4 | 0.04 | 0.00 | 0.04 | 0.01 | 0.56 | 0.01 |
| 2015 Winter | ST 1 | 0.04 | 0.01 | 0.30 | 0.01 | 0.86 | 0.03 |
|  | ST 1A | 0.05 | 0.00 | 0.16 | 0.00 | 0.77 | 0.00 |
|  | ST 2 | 0.04 | 0.00 | 0.28 | 0.12 | 0.66 | 0.00 |
|  | ST 2B | 0.03 | 0.00 | 0.19 | 0.04 | 0.55 | 0.03 |
|  | ST 3 | 0.02 | 0.00 | 0.11 | 0.01 | 0.62 | 0.02 |
|  | ST 4 | 0.02 | 0.00 | 0.23 | 0.02 | 0.60 | 0.00 |
| 2016 Spring | ST 1 | 0.04 | 0.00 | 0.05 | 0.01 | 0.86 | 0.01 |
|  | ST 1A | 0.04 | 0.00 | 0.04 | 0.01 | 0.60 | 0.02 |
|  | ST 2 | 0.04 | 0.01 | 0.04 | 0.00 | 0.59 | 0.07 |
|  | ST 2B | 0.03 | 0.00 | 0.06 | 0.01 | 0.68 | 0.01 |
|  | ST 3 | 0.03 | 0.00 | 0.48 | 0.04 | 0.83 | 0.04 |
|  | ST 4 | 0.03 | 0.01 | 0.22 | 0.02 | 0.87 | 0.00 |
| 2016 Summer | ST 1 | 0.03 | 0.00 | 0.03 | 0.01 | 1.00 | 0.11 |
|  | ST 1A | 0.02 | 0.01 | 0.02 | 0.01 | 0.64 | 0.02 |
|  | ST 2 | 0.01 | 0.01 | 0.04 | 0.00 | 0.64 | 0.04 |
|  | ST 2B | 0.02 | 0.00 | 0.03 | 0.00 | 0.68 | 0.01 |
|  | ST 3 | 0.01 | 0.01 | 0.03 | 0.00 | 0.67 | 0.03 |
|  | ST 4 | 0.02 | 0.01 | 0.04 | 0.01 | 0.71 | 0.02 |

Supplementary Table S3: Nutrient data from fresh unfrozen samples of the July 2016 cruise. *acidified samples were unfrozen 5 days before analysis. All ammonium and phosphate concentrations are below effective detection limits.

| Cruise | Station | Phosphate (nM) | Nitrate (nM) | Ammonia (nM) | Silicate (µM) * |
| --- | --- | --- | --- | --- | --- |
| Summer  2016  (fresh samples) | ST 1 | 5.1 | 43.7 | 4.2 | 1.52 |
|  | ST 1A | 0.2 | 20 | 1.5 | 0.92 |
|  | ST 2 | 0.7 | 15.5 | 1.3 | 0.85 |
|  | ST 2B | 2.1 | 28.8 | 6.1 | 0.8 |
|  | ST 3 | 1.4 | 17.3 | 5 | 0.83 |
|  | ST 4 | 1.4 | 8.4 | 7 | 0.75 |

Supplementary Table S4: Pigment concentrations in ng/l based on UPLC measurements.

Abbreviations for pigments: Chl *a*: chlorophyll *a*, 19-Hx: 19’-hexanoyloxyfucoxanthin, 19-Bx: 19’-butanoyloxyfucoxanthin, Fuco: fucoxanthin, Per: peridinin, DV-Chl *a*: divinyl chlorophyll *a*, Chl C2a: chlorophyll *c* peak 1, Chl C2b: chlorophyll *c* peak 2 (Chl C2 was separated as two peak due to running conditions), Chl *b*: chlorophyll *b*, b-Car: beta-carotene, Zea: zeaxanthin, Diadino: diadinoxanthin, Diato: diatoxanthin, Dino: dinoxanthin.

Other abbreviations: - : no sample, ND: not detected

| **Year Season** | **Station** | **Chl *a*** | **19-Hx** | **19-Bx** | **Fuco** | **Per** | **DV-Chl *a*** | **Chl C2a** | **Chl C2b** | **Chl *b*** | **b-Car** | **Zea** | **Diadino** | **Diato** | **Dino** |
| --- | --- | --- | --- | --- | --- | --- | --- | --- | --- | --- | --- | --- | --- | --- | --- |
| 2014 Winter | ST 1 | - | - | - | - | - | - | - | - | - | - | - | - | - | - |
|  | ST 2 | 128.53 | 78.96 | 36.45 | 13.89 | ND | ND | 8.93 | 12.65 | ND | ND | 28.00 | 3.54 | ND | 0.75 |
|  | ST 3 | 44.04 | 55.94 | 23.04 | 6.57 | ND | ND | 15.49 | ND | ND | 2.98 | 12.67 | 1.70 | ND | ND |
|  | ST 4 | 39.94 | 46.43 | 22.98 | 7.98 | ND | ND | 4.89 | 4.36 | ND | ND | 18.55 | 1.98 | ND | 0.44 |
| 2015 Spring | ST 1 | 57.31 | 24.52 | 10.63 | 7.22 | ND | ND | 4.28 | 4.43 | ND | ND | 9.97 | 2.69 | ND | 0.66 |
|  | ST 2 | 47.79 | 35.19 | 15.33 | 8.94 | ND | 0.44 | 9.13 | 10.33 | 0.86 | ND | 11.49 | 3.31 | 0.57 | 0.93 |
|  | ST 3 | 47.80 | 18.16 | 7.18 | 3.19 | ND | ND | 4.70 | 8.66 | ND | ND | 7.22 | 2.44 | 0.41 | 0.67 |
|  | ST 4 | 58.37 | 25.76 | 10.07 | 6.86 | 2.78 | ND | 4.87 | 7.99 | ND | ND | 13.98 | 2.72 | ND | 0.97 |
| 2015 Summer | ST 1 | 88.59 | 34.81 | 8.43 | 5.93 | ND | ND | 4.39 | 7.81 | ND | ND | 24.68 | 3.40 | 0.63 | 0.87 |
|  | ST 2 | - | - | - | - | - | - | - | - | - | - | - | - | - | - |
|  | ST 3 | 37.40 | 25.36 | 3.53 | 2.65 | 1.56 | ND | 2.24 | 2.22 | ND | ND | 7.59 | 2.10 | 1.14 | 0.84 |
|  | ST 4 | 58.53 | 29.28 | 4.17 | 3.36 | 2.94 | ND | 1.92 | 2.36 | 1.20 | ND | 8.99 | 2.94 | 1.77 | 1.12 |

| **Year Season** | **Station** | **Chl *a*** | **19-Hx** | **19-Bx** | **Fuco** | **Per** | **DV-Chl *a*** | **Chl C2a** | **Chl C2b** | **Chl *b*** | **b-Car** | **Zea** | **Diadino** | **Diato** | **Dino** |
| --- | --- | --- | --- | --- | --- | --- | --- | --- | --- | --- | --- | --- | --- | --- | --- |
| 2015 Winter | ST 1 | - | - | - | - | - | - | - | - | - | - | - | - | - | - |
|  | ST 2 | 160.86 | 78.67 | 18.57 | 13.39 | ND | ND | 10.00 | 9.27 | ND | ND | 12.58 | 5.72 | ND | ND |
|  | ST 3 | 153.61 | 101.34 | 21.85 | 10.80 | ND | ND | 11.83 | 11.92 | ND | ND | 14.67 | 7.52 | ND | 3.57 |
|  | ST 4 | 473.63 | 109.38 | 25.16 | 11.48 | ND | ND | 12.95 | 13.75 | ND | 2.08 | 28.85 | 13.28 | ND | 7.72 |
| 2016 Spring | ST 1 | 127.53 | 19.95 | 8.09 | 8.36 | 5.08 | 1.86 | 2.40 | 3.18 | 1.38 | ND | 22.26 | ND | 1.18 | ND |
|  | ST 2 | 84.22 | 9.95 | 2.38 | 1.12 | ND | 2.04 | ND | 0.80 | 0.94 | 0.60 | 15.27 | 2.92 | 1.65 | ND |
|  | ST 3 | 77.29 | 16.81 | 5.84 | 3.65 | 2.27 | 1.26 | ND | 0.96 | 3.73 | 0.33 | 22.81 | 4.49 | 1.53 | ND |
|  | ST 4 | 179.22 | 18.42 | 6.86 | 6.76 | 2.81 | 9.06 | ND | 2.47 | 2.71 | 0.69 | 36.29 | 4.78 | 1.79 | 0.65 |
| 2016 Summer | ST 1 | 156.92 | 15.41 | 2.10 | 3.19 | 0.75 | 0.54 | 0.81 | 2.32 | 0.99 | ND | 37.00 | 7.81 | 0.64 | ND |
|  | ST 2 | 55.47 | 5.58 | 0.94 | 0.86 | 0.20 | ND | 0.28 | 0.67 | ND | ND | 10.78 | 3.99 | 0.56 | ND |
|  | ST 3 | 129.44 | 8.93 | 1.57 | 1.27 | ND | ND | 0.56 | 1.46 | 1.70 | 0.92 | 21.08 | 6.16 | 1.02 | ND |
|  | ST 4 | 177.49 | 5.20 | 0.98 | 1.20 | 1.67 | ND | 0.66 | 1.47 | 4.43 | ND | 22.90 | 6.52 | 0.80 | 2.21 |

Supplementary Table S5: CTD data at 10m for the stations sampled for DNA and RNA. Due to technical problems, no salinity value was obtained for station 1 in the Spring 2016 cruise. For the environmental matrix, the value from station 1A was used and is given in brackets.

| Year  Season | Station | Temperature [°C] | Salinity [PSU] | Fluorescence | Turbidity [FTU] |
| --- | --- | --- | --- | --- | --- |
| 2014  early  Winter | ST 1 | 22.57 | 39.32 | 0.140 | 0.151 |
|  | ST 2 | 22.43 | 39.34 | 0.108 | 0.149 |
|  | ST 3 | 22.47 | 39.31 | 0.096 | 0.131 |
|  | ST 4 | 22.43 | 39.35 | 0.117 | 0.131 |
| 2015 Spring | ST 1 | 17.86 | 38.82 | 0.068 | 0.161 |
|  | ST 2 | 17.73 | 38.83 | 0.061 | 0.117 |
|  | ST 3 | 17.64 | 38.81 | 0.053 | 0.066 |
|  | ST 4 | 17.72 | 38.87 | 0.083 | 0.085 |
| 2015 Summer | ST 1 | 26.22 | 39.04 | 0.063 | 0.832 |
|  | ST 2 | 26.68 | 39.24 | 0.052 | 1.341 |
|  | ST 3 | 26.52 | 39.27 | 0.042 | 1.022 |
|  | ST 4 | 26.59 | 39.24 | 0.045 | 1.390 |
| 2015  early  Winter | ST 1 | 24.37 | 39.38 | 0.509 | 0.208 |
|  | ST 2 | 24.27 | 39.26 | 0.436 | 0.094 |
|  | ST 3 | 24.04 | 39.21 | 0.400 | 0.090 |
|  | ST 4 | 24.08 | 39.24 | 0.277 | 0.073 |
| 2016 Spring | ST 1 | ­18.59 | (38.88) | 0.136 | 0.206 |
|  | ST 2 | 18.24 | 38.93 | 0.067 | 0.143 |
|  | ST 3 | 18.22 | 38.93 | 0.068 | 0.117 |
|  | ST 4 | 18.44 | 38.91 | 0.092 | 0.215 |
| 2016 Summer | ST 1 | 28.43 | 39.17 | 0.076 | 6.219 |
|  | ST 2 | 28.40 | 39.47 | 0.051 | 3.895 |
|  | ST 3 | 28.42 | 39.46 | 0.133 | 3.897 |
|  | ST 4 | 28.66 | 39.39 | 0.075 | 3.967 |

Supplementary Table S6: Flow cytometry data in cells/ml. Samples for *Prochlorococcus*, *Synechococcus*, Picoeurkaryotes and 'Total cells (BD)' were all run on a BD cantor flow cytometer. A Guava Easycyte flow cytometer was used for 'Total cells (Guava)'. Samples for total cell counts were stained with Sybr Green. Samples for *Prochlorococcus*, *Synechococcus* and Picoeurkaryotes were not stained and counted base on autofluorescence signals. Samples were run in triplicates and for technical replicates were run for each sample on the BD cantor. ND: not detected

| Year  Season | Station | *Prochloroccous* | | *Synechococcus* | | Picoeukaryotes | | Total cells (BD) | | Total cells (Guava) | |
| --- | --- | --- | --- | --- | --- | --- | --- | --- | --- | --- | --- |
|  |  | Average | SD | Average | SD | Average | SD | Average | SD | Average | SD |
| 2014 early  Winter | ST 1 | 4223 | 102 | 4800 | 266 | 39091 | 4052 | 538408 | 85443 | 1234031 | 32480 |
|  | ST 2 | 9022 | 464 | 10978 | 632 | 4106 | 2395 | 619542 | 80394 | 803126 | 157212 |
|  | ST 3 | 10303 | 1136 | 14330 | 792 | 6109 | 765 | 634490 | 112194 | 677237 | 187557 |
|  | ST 4 | 11486 | 1188 | 9824 | 571 | 5445 | 2836 | 939931 | 435038 | 1068824 | 209741 |
| 2015 Spring | ST 1 | 15966 | 1777 | 9302 | 1116 | ND |  | 897962 | 420694 | 1052595 | 323774 |
|  | ST 2 | 18913 | 410 | 8534 | 880 | ND |  | 1008005 | 218981 | 922332 | 198639 |
|  | ST 3 | 3498 | 554 | 8717 | 328 | ND |  | 977237 | 162732 | 935803 | 109145 |
|  | ST 4 | 7477 | 1137 | 9904 | 1585 | ND |  | 481180 | 34865 | 736400 | 37796 |
| 2015 Summer | ST 1 | ND |  | 8818 | 2666 | ND |  | 259249 | 23515 | 507848 | 20230 |
|  | ST 2 | ND |  | 3187 | 131 | ND |  | 207478 | 10469 | 310475 | 8128 |
|  | ST 3 | ND |  | 8259 | 1838 | ND |  | 234948 | 38054 | 336711 | 7070 |
|  | ST 4 | ND |  | 5983 | 889 | 13939 | 7981 | 176812 | 19239 | 310250 | 9665 |
| 2015 early Winter | ST 1 | 5319 | 662 | 14266 | 346 | 4021 | 211 | 459085 | 144217 | 1090478 | 33143 |
|  | ST 2 | 5262 | 737 | 16589 | 6296 | 5496 | 269 | 417091 | 21952 | 603783 | 41544 |
|  | ST 3 | 2851 | 1026 | 14979 | 6569 | 4000 | 1366 | 325249 | 52443 | 375638 | 7672 |
|  | ST 4 | 3468 | 334 | 13837 | 5596 | 4362 | 391 | 295332 | 138239 | 358828 | 16121 |
| 2016 Spring | ST 1 | 26182 | 1639 | 19494 | 1013 | ND |  | 466767 | 54832 | 714629 | 30020 |
|  | ST 2 | 12092 | 674 | 15880 | 681 | ND |  | 576033 | 26706 | 592979 | 16383 |
|  | ST 3 | 28175 | 23272 | 9165 | 1382 | ND |  | 506129 | 18949 | 576112 | 29628 |
|  | ST 4 | 36318 | 517 | 20475 | 1291 | ND |  | 562133 | 13642 | 709786 | 30788 |
| 2016 Summer | ST 1 | 31546 | 6119 | 13946 | 486 | 15348 | 6312 | 1043058 | 182054 | 793743 | 36561 |
|  | ST 2 | 9736 | 346 | 8194 | 235 | 1927 | 206 | 397337 | 8553 | 381659 | 6573 |
|  | ST 3 | 9271 | 416 | 7854 | 224 | 2723 | 957 | 419149 | 48082 | 362844 | 14128 |
|  | ST 4 | 18942 | 6726 | 6509 | 560 | 8120 | 3172 | 233578 | 18835 | 424307 | 11646 |

Supplementary Table S7: Relation between abundance in RNA and DNA samples for families contributing >= 1% of total reads. First correlation between abundance in DNA and RNA samples is given followed by the family level summary of the DEseq2 analysis. Significant P-values are in bold.

| Taxonomy | Average RNA:DNA ratio  ± standard deviation | Spearman rs | *P*-value | DEseq2 results | | |
| --- | --- | --- | --- | --- | --- | --- |
|  |  |  |  | OTUs sign. up in RNA | OTUs sign. up in DNA | Sign. OTUs % of RNA reads / % of DNA reads |
| Proteobacteria; Rhodobacterales; Rhodobacteraceae | 1.45 ± 0.72 | 0.0531 | 0.788 | 14 | 0 | 2.2 / 1.4 |
| Actinobacteria; Acidomicrobiales; OM1_clade | 0.02 ±0.02 | 0.1759 | 0.371 | 0 | 17 | 0.04 / 2.5 |
| Proteobacteria; Rhodospirillales; Rhodospirillaceae | 2.66 ± 0.85 | 0.2633 | 0.176 | 84 | 3 | 6.7 / 2.1 |
| Bacteriodetes; Flavobacteriales; Flavobacteriaceae | 0.19 ± 0.07 | 0.4428 | **0.018** | 0 | 64 | 1.3 / 7.0 |
| Cyanobacteria; Subsection I;  Family I | 2.44 ± 0.86 | 0.7083 | **<0.001** | 150 | 0 | 23.5 / 9.7 |
| Marinimicrobia SAR406 clade | 0.76 ± 0.41 | 0.9407 | **<0.001** | 14 | 7 | 1.3 / 1.6 |
| Proteobacteria; Oceanospirillales; SAR86 clade | 1.13 ± 0.27 | 0.7154 | **<0.001** | 14 | 0 | 2.9 / 2.4 |
| Proteobacteria; Rickettsiales;  S25-593 | 1.34 ± 0.35 | 0.6538 | **<0.001** | 5 | 0 | 0.8 / 0.5 |
| Proteobacteria; Rickettsiales; SAR116_clade | 1.74 ± 0.48 | 0.8528 | **<0.001** | 44 | 0 | 7.4 / 3.8 |
| Proteobacteria; SAR11 clade I | 0.34 ± 0.09 | 0.6716 | **<0.001** | 0 | 211 | 10.9 / 32.2 |
| Proteobacteria; SAR11 clade II | 0.15 ± 0.04 | 0.6954 | **<0.001** | 0 | 45 | 0.8 / 5.6 |
| Proteobacteria; SAR11 clade IV | 0.84 ± 0.35 | 0.9501 | **<0.001** | 1 | 0 | 0.0039 / 0.0004 |
| Verrucomicrobia; Puniceicoccales; Puniceicoccaceae | 1.21 ± 0.37 | 0.8496 | **<0.001** | 2 | 0 | 0.06 / 0.04 |

Supplementary Table S8: Statistics of permutational multivariate analysis of variance using Bray-Curtis distances and 1000 permutations performed using Adonis of the R package vegan.

| Sample type |  | Df | Sums of Sqaures | Mean Squares | F.Model | R^2^ | Pr(>f) |
| --- | --- | --- | --- | --- | --- | --- | --- |
| RNA | Season | 2 | 1.8402 | 0.92008 | 18.0167 | 0.53318 | 0.000999 |
|  | Station | 1 | 0.2095 | 0.20953 | 4.1029 | 0.06071 | 0.003996 |
|  | Season;Station | 2 | 0.2781 | 0.13904 | 2.7227 | 0.08058 | 0.004995 |
|  | Residual | 22 | 1.1235 | 0.05107 |  | 0.32553 |  |
|  | Total | 27 | 3.4513 |  |  | 1.00000 |  |
| DNA | Season | 2 | 0.97998 | 0.48999 | 15.2324 | 0.49800 | 0.000999 |
|  | Station | 1 | 0.11484 | 0.11484 | 3.5701 | 0.05836 | 0.003996 |
|  | Season;Station | 2 | 0.16531 | 0.08266 | 2.5696 | 0.08401 | 0.004995 |
|  | Residual | 22 | 0.70769 | 0.03217 |  | 0.35963 |  |
|  | Total | 27 | 1.96783 |  |  | 1.00000 |  |

Supplementary Table S9: Overlap of clustering of OTUs with more than 0.1% of total reads in DNA and/or RNA samples. Colored cells indicated OTUs clustered in both analyses. Color indicates seasonal preference of the cluster: blue – early winter, green – spring, orange – summer, no seasonal preference - grey. Unclustered: OTUs with >0.1% of total reads in DNA and RNA but without assignment to a cluster in one of the analysis. #N/A: OTUs with less than 0.1% total reads in either DNA or RNA.

| **RNA\DNA cluster** | **D1** | **D2** | **D3** | **D4** | **D5** | **D6** | **D7** | **D8** | **D9** | **D10** | **Unclustered**  **DNA** | **#N/A DNA** | **Number of OTUs in RNA cluster** |
| --- | --- | --- | --- | --- | --- | --- | --- | --- | --- | --- | --- | --- | --- |
| **R1** |  |  |  |  |  |  | 5 |  |  |  |  | 1 | 6 |
| **R2** |  | 4 | 1 |  |  |  |  |  |  |  | 1 | 7 | 13 |
| **R3** |  |  |  | 1 | 2 |  |  | 1 |  |  |  | 4 | 8 |
| **R4** |  |  |  |  |  | 10 | 1 |  |  |  |  | 2 | 13 |
| **R5** |  |  |  |  |  |  |  |  |  |  |  | 5 | 5 |
| **R6** |  |  |  |  |  |  |  |  |  |  |  | 4 | 4 |
| **R7** |  |  |  |  |  |  |  |  |  | 3 |  | 13 | 16 |
| **R8** |  |  | 1 |  |  |  |  |  |  | 2 | 1 | 4 | 8 |
| **R9** | 5 | 1 |  | 2 | 12 |  |  | 4 |  |  | 2 | 5 | 31 |
| **R10** |  |  |  |  |  | 1 |  |  | 5 | 1 |  | 4 | 11 |
| **Unclustered RNA** | 1 |  | 2 | 2 | 1 | 1 |  |  | 1 | 2 | 1 | 5 | 16 |
| **#N/A RNA** | 2 | 1 | 3 |  | 2 | 7 | 2 | 2 | 4 | 3 | 7 |  | 33 |
| **Number of OTUs in DNA cluster** | 8 | 6 | 7 | 5 | 17 | 19 | 8 | 7 | 10 | 11 | 12 | 54 |  |
