## Supplementary Information for "Microbial communities in an ultra-oligotrophic sea are more affected by season than by distance from shore"

For the article:

Authors’ affiliations:

^1^ Marine Microbiology Lab, Department of Marine Biology, Leon H. Charney School of Marine Sciences, Haifa University

^2^ Department of Aquatic Microbial Ecology, Institute of Hydrobiology, Biology Centre CAS, Czech Republic

^3^ Bioinformatics Service Unit, University of Haifa

^4^ The Dr. Moses Strauss Department of Marine Geosciences, Leon H. Charney School of Marine Sciences, Haifa University

^5^ Morris Kahn Marine Research Station, Environmental Geochemistry Lab., Leon H. Charney School of Marine Sciences, Haifa University

Content	Page

Supplementary Information on Methods

Nutrient analysis	2

Flow cytometry	2

Pigment analysis	3

DNA and RNA extraction	4

PCR amplification and sequencing of 16S rRNA genes (DNA)

and transcripts (RNA) samples	5

Sequence processing	6

Analysis	7

Supplementary Information on Results

Phytoplankton structure	8

References	10

**Supplementary Information on Methods**

*Nutrient analysis*

Silicate, nitrate+nitrite and soluble reactive phosphate of frozen samples were measured calorimetrically on a LACHAT Instruments Quik-Chem 8500 Flow Injection Analyzer (Grasshoff *et al.*, 2009) at the service unit of the Interuniversity Institute for Marine Sciences in Eilat, Israel. The internal precision (the average deviation from the mean of duplicate samples) of the measurements and the effective limit of detection for these frozen samples were 0.04 μmol/L for phosphate and 0.05 μmol/L for both silicate and nitrate+nitrite, respectively. Accuracy was obtained by calibration against a suite of solutions prepared by dilution of commercial, high-concentration standards (1000 mg/L, Merck). The calibration curve was run several times during every flow injection analyzer session.

During the July 2016 cruise nutrient concentrations were also determined from non-frozen samples. These samples were analysed within 24 hours of collection at the Morris Kahn Marine Laboratory, Sdot Yam, Israel, using a SEAL AA-3 autoanalyzer. Silicate, nitrate+nitrite and soluble reactive phosphate were measured calorimetrically following the manufacturer’s operating protocols (SEAL version 2015). Silicate was measured by a modification of the silico-molybdate blue method (Grasshoff *et al.*, 1983), nitrate+nitrite determined by a purple diazo dye after Cd reduction (Armstrong *et al*. 1967), while soluble reactive phosphate was determined using a long flow cell (50 cm) and a modified Murphy and Riley (1962) molybdophosphate blue method. Ammonia was determined using a fluorometric method based on (Kérouel and Aminot, 1997). The precision of these methods at the range used for the Eastern Mediterranean Sea surface waters were 33 nM, 21 nM, 2 nM and 6 nM, respectively, and the limits of detection were 55nM, 63nM, 2nM and 5 nM (defined as 3 times the precision of the seawater blank).

*Flow cytometry*

To each sample run on the BD flow cytometer, 2 μm diameter fluorescent beads (Polysciences, Warminster, PA, USA) were added as an internal standard. At first, the natural fluorescence of the cells (chlorophyll and phycoerythrin pigments) was examined to identify three cell types: i) *Prochlorococcus* and ii) picoeukaryotes, which were identified based on the PerCP channel and forward scatter with picoeukaroytes being larger and having higher fluorescence than *Prochlorococcus*); iii) *Synechococcus* cells, which were identified based on their emission in the PE channel. Samples were then stained with SYBR Green I (Molecular Probes/ ThermoFisher) to a final concentration according to the manufacturer’s instructions and the total microbial population was counted. Data were acquired and processed with FlowJo software. Flow rates were determined several times during each running session, and the average value for a specific run was used for calculating cells per ml.

For runs on the Guava easyCyte HT, 200 µl of each sample was transferred into 96 well plates, stained for 45 min in the dark with 2 µl of 1:20 in TE diluted SYBR Green I and 4-fold serial diluted (up to 1024x) with TE. Samples were run and analyzed using the Guava Express Plus suite. Dilutions with 100 to 500 cells/µl were used in the analysis. SAR11 HTCC1062 cells were run in parallel as standard.

*Pigment analysis*

The protocol for pigment analysis was based on LOV method (Hooker et al. 2005) with some adjustments to UPLC. The biomass collected on glass fiber filters was extracted with 1 ml of 100% methanol for 2.5 h at room temperature. Obtained organic extracts were immediately filtered through syringe filters (Acrodisc CR, 13 mm, 0.2 µm PTFE membranes, Pall Life Sciences) and transferred to UPLC vials. 10µl of preheated (30°C) extract analyzed using an ACQUITY UPLC system (Waters) equipped with a a PDAe λ detector (Waters). Separation of extracts was performed on a reverse phase C8 column (1.7 µm particle size, 2.1 mm internal diameter, 50 mm column length, ACQUITY UPLC BEH, 186002877) heated to 50°C. Flow rate was set 0.5 mL/min and total runtime to 14 min. The mobile phase consistent of a gradient of solvents A (70:30 mixture of methanol and 0.5 M ammonium acetate) and B (100% methanol). The gradient was as follows: 80/20 solvent A/B from 0 to 0.2 min, linear decrease to 50/50 solvent A/B at 2 min and to 0/100 solvent A/B at 9 min and kept at there until 12 min. The column was reset by increasing the gradient linear to 80/20 solvent A/B at 12.5 min and keeping it there till the end of the run at 14 min. Peaks were monitored at 440 nm and identified by comparing its retention time and spectrum absorbance (obtained by PDA detector reads at 350-700 nm) against the known standards of chlorophyll *a*, divinyl-chlorophyll *a*, chlorophyll *b*, chlorophyll *c*2, zeaxanthin, betacarotene, diatoxanthin, dinoxanthin, fucoxanthin and peridinin, which were analyzed before each run to ease identification and calculate pigment concentrations. All standards were purchased from the DHI, Denmark, except chlorophyll *a*, which was purchased from Sigma. The fucoxanthin standard was also used to estimate concentrations of 19’-hexanoylfucoxanthin and 19’-butanoylfucoxanthin. Due to problems with the stability of the chlorophyll *a* standard, chlorophyll *a* concentrations were evaluated by the known factor of 3.5 between chlorophyll *b* to chlorophyll *a* µg/unit area (Rpt) that was calibrated previously using the same separation method (data not shown).

*DNA and RNA extraction*

Nucleic acids were extracted at the BioRap unit, Faculty of Medicine, Technion, Israel using a semi-automated protocol, which includes manually performed chemical and mechanical cell lysis before the automated steps. The manual part began with thawing the samples. From the sterivex filters, the storage buffer was removed using a syringe and 170 µl lysis buffer (20 mM Tris HCl pH 8.0, 2 mM EDTA, 1.2% Triton) was added to the filter. RNA Save was completely removed from RNA filters and 170 µl lysis buffer were added. Bead-beating was carried out at 30 Hz for 1.5 min using the TissueLyser LT^TM^ (Qiagen) with two 3 mm stainless steel balls. After addition of 30 µl lysozyme (20 mg/ml), RNA and DNA samples were incubated at 37°C for 30 min. 20 µl proteinase K and 200 µl AL buffer (Qiagen) were added and the tubes and sterivex filters incubated for 1 hour at 56°C with agitation. The liquid part of the sterivex filters was eluted into tubes. Both RNA and DNA samples were then centrifuged for 10 min at 5000 x g and the supernatant transferred to a new tube, which was subjected to the QIAcube automated system (Qiagen). DNA was extracted following the manufacturer's instructions using the QIAamp DNA Mini Protocol: DNA Purification from Blood or Body Fluids (Spin Protocol) from step 6 and onwards. All DNA samples were eluted in 100 μl DNA free distilled-water. RNA was extracted using the protocol for purification of total RNA from bacteria with the RNeasy Mini Kit, which includes a DNase treatment step. RNA samples were eluted with 30 µl RNase-free water. Extracted DNA and RNA samples were quality checked on a tape station and quantified using the PicoGreen assay.

### *PCR amplification and sequencing of 16S rRNA genes (DNA) and transcripts (RNA) samples*

### A two-stage “targeted amplicon sequencing” protocol (*e.g.* (Bybee *et al.*, 2011; Green *et al.*, 2015)) was performed to PCR amplify the 16S rRNA gene from cDNA and DNA. The primers used in the first PCR stage consisted of the 16S primer set 515F-Y and 926R (Parada *et al.*, 2016) that targets the variable V4-5 region with common sequence tags (CS1 and CS2) added at the 5’ end as described previously (*e.g.* (Moonsamy *et al.*, 2013)). The first PCR stage was performed in triplicates in a total volume of 25 μl containing 0.5 ng of template, 12.5 μl of MyTaq Red Mix (Bioline), 0.5 μl of 10 μM forward CS1_515F-Y (ACACTGACGACATGGTTCTACAGTGYCAGCMGCCGCGGTAA) and reverse CS2_926R (TACGGTAGCAGAGACTTGGTCTCCGYCAATTYMTTTRAGTTT) primers. Amplification was performed by an initial denaturation step at 95°C for 5 min, 28 cycles at 95°C for 30 sec, 50°C for 30 sec and 72°C for 1 min followed by a final elongation step for 5 min at 72°C. All PCR products were validated on 1% agarose gels and then triplicates were pooled. All RNA samples were tested for the presence of contaminating DNA in the RNA samples by PCR on the RNA samples without the reverse transcription step.

Subsequently, a second PCR amplification was performed in 10 μl reactions in 96-well plates. A mastermix for the entire plate was made using the MyTaq HS 2X mastermix. Each well received a separate primer pair with a unique 10-base barcode, obtained from the Access Array Barcode Library for Illumina (Fluidigm, South San Francisco, CA; Item# 100-4876). These AccessArray primers contained the CS1 and CS2 linkers at the 3’ ends of the oligonucleotides. Cycling conditions were as follows: 95°C for 5 min, followed by 8 cycles of 95°C for 30 sec, 60°C for 30 sec and 72°C for 30 sec and a final elongation at 72°C for 7 min. Samples were pooled in equal volume using an EpMotion5075 liquid handling robot (Eppendorf, Hamburg, Germany). The pooled library was purified using an AMPure XP cleanup protocol (0.6X, vol/vol; Agencourt, Beckmann-Coulter) to remove fragments smaller than 300 bp. The pooled libraries, with a 20% phiX DNA spike-in, were loaded onto an Illumina MiniSeq mid-output flow cell (2x150 paired-end reads). Based on the distribution of reads per barcode, the amplicons (before purification) were re-pooled to generate a more balanced distribution of reads. The re-pooled library was purified using AMPure XP cleanup, as described above. The re-pooled libraries, with a 15% phiX DNA spike-in, were loaded onto a MiSeq v2 flow cell, and sequenced (2x250 paired-end reads) using an Illumina MiSeq sequencer. Fluidigm sequencing primers, targeting the CS1 and CS2 linker regions, were used to initiate sequencing. De-multiplexing of reads was performed on instrument. Library preparation, pooling, and MiniSeq sequencing were performed at the DNA Services (DNAS) facility, Research Resources Center (RRC), University of Illinois at Chicago (UIC). MiSeq sequencing was performed at the W.M. Keck Center for Comparative and Functional Genomics at the University of Illinois at Urbana-Champaign (UIUC).

*Sequence processing*

For each sample the obtained pair of fastq files were subjected to quality control prior to analysis. QC steps were carried out with MOTHUR Version 1.40.2 (Schloss *et al.*, 2009). First, paired reads were assembled into contigs using the make.contigs command with default parameters, except for deltaq=4 and insert=30. Then, sequences with any ambiguity, homopolymers longer than 8 nt and sequences shorter than 405 or longer than 420 nt were filtered out. Suspected chimera were detected using the chimera.uchime algorithm using the Silva.gold.align database as reference, and removed. The filtered dataset was clustered into operational taxonomic units (OTUs) using QIIME pick_otus.py script with default parameters. Similarity level was set at 97% threshold. Representative sequences for each OTU were obtained by qiime pick_rep_set.py script with default parameters. OTUs were then classified using MOTHUR classify.seqs command with silva.nr_v128.align as template. Confidence level was set to 80%. OTUs for which total sequence counts across all samples were <10 sequences were removed. In addition, OTUs identified of non-prokaryotic, chloroplasts- or mitochondria-origin were removed. In total, the 56 samples had 2,820,858 quality sequences, which binned into 9448 OTUs (at 97% similarity threshold). In order to avoid bias related to differences in library size, all libraries were rarified to 20,000 reads per sample, using the rrarify command implemented in the R package Vegan (Oksanen et al. 2017).

*Analysis*

The R package Vegan was used for non-metric multidimensional scaling analysis based on the Bray-Curtis dissimilarity matrix with the following parameters: K=2, 100 runs and 100 iterations. The Vegan envfit command with 999 permutations was used to calculate Goodness of fit between different sample characteristic and environmental factors. The Vegan ‘adonis’ command was used based on Bray-Curtis dissimilarities with 999 permutations in order to examine significance of grouping factors, such as molecule type, season and sampling station, the ADONIS test was applied. Where required, post-hoc pairwise Adonis test were calculated using the pairwise.adonis command.

Correspondence between abiotic and biotic measurements and variation in community composition was examined by variation partitioning analysis and canonical correspondence analysis. First, the explanatory variables proposed were divided into three matrices: I) Physical: distance from shore, temperature, salinity, turbidity; II) Nutrients: phosphate, NO_3_+NO_2_, silicate; III) Biotic: total fluorescence, total cell counts, *Prochlorococcus* cell counts, *Synechococcus* cell counts. Variation partitioning calculates the canonical coefficient of determination (adj. *R^2^*) representing the forecasting potential of multiple regression between the response variables (*i.e*., the OTUs) and explanatory variables (*e.g*., environmental measurements). Variation partitioning was performed using the R package Vegan ‘varpart’ command. Permutation tests, to validate the significance of the coefficients (adj. R^2^) were performed both directly and conditionally (excluding variation that is related to interactions between the different explanatory variable matrices) by calculating canonical correspondence analysis (CCA) using the ‘cca’ command followed by ‘anova.cca’ for the model, the tested variables and the CCA axes.

**Supplementary Information on Results**

*Phytoplankton structure*

UPLC-based pigment analysis and flow cytometry suggested seasonal changes in phytoplankton abundance and community structure. The most abundant and commonly found pigments apart from chlorophyll *a* were in order of average concentrations: 19’-hexanoyloxyfucoxanthin (19-Hx), zeaxanthin, 19’-butanoyloxyfucoxanthin (19-Bx), and fucoxanthin (Supplementary Table S4). 19-Hx concentrations peaked in winter samples with much lower concentrations detected in spring and summer (Supplementary Figure S1 A). 19-Bx and fucoxanthin showed the similar patterns (Supplementary Figure S1 B, C). In winter samples, 19-Hx was 5.7-9.5 times and 19-Bx 1.4-3.5 times more abundant than fucoxanthin. The observed ratios indicate a potential bloom of pigment type 8 haptophytes *sensu* (Zapata *et al.*, 2004) as this is the only haptophyte pigment type with 19-Hx as the main pigment, 19-Bx in more than trace amounts, and a ratio of 19-Bx to fucoxanthin above 1. Pigment type 8 has been found in the haptophyte family Phaeocystaceae and members of the Prymnesiaceae and Isochrysidaceae families (Zapata *et al.*, 2004). Dinoflagellate or diatom blooms seemed less likely for several reasons. Dinoflagellates of pigment type 2 and 3 *sensu* (Zapata *et al.*, 2012) have a combination of 19-Hx, 19-Bx and fucoxanthin, however when 19-Hx is the dominant pigment, fucoxanthin is usually much more abundant than 19-Bx (Zapata *et al.*, 2012). In diatoms fucoxanthin is the main photosynthetic carotenoid (Kuczynska *et al.*, 2015), but 19-Bx has only been occasionally reported while 19-Hx has not been detected (Jeffrey *et al.*, 2011). While 19-Hx and 19-Bx showed clear concentration peaks in winter and very low concentrations in both spring and summer, fucoxanthin concentrations decreased more gradually from winter to summer and concentration remained relatively high at station 1 (Supplementary Figure S1 C), indicating the potential additional presence of diatoms especially at station 1 in both studied springs and summer 2015.

Dinoflagellates of pigment type 1 *sensu* (Zapata *et al.*, 2012), easily characterized by the specific pigment peridinin, were absent in winter samples at all stations, but present in all spring and summer samples of station 4 and some samples of station 1, 2 and 3 (Supplementary Table S4).

Divinyl-chlorophyll *a*, a characteristic pigment of *Prochlorococcus* Cyanobacteria, was not detected in the winter cruises nor in the summer cruises, with the exception of station 1 in summer 2016 (Supplementary Table S4). During spring 2015 it was detected in station 2 and in all four stations in spring 2016 with the highest amounts in the most offshore station 4. This pattern was similar to *Prochlorococcus* abundance based on flow cytometry data (Supplementary Figure S5 B).

*Synechococcus* cyanobacteria showed small seasonal difference in abundance at stations 2, 3 and 4, as detected by flow cytometry (Supplementary Figure S5 A). Within each cycle of cruises (winter, spring, summer), *Synechococcocus* abundances tended to be lowest in summer. The same trend was observed in zeaxanthin abundance (Supplementary Figure S1 D), which has been used as signature pigment for cyanobacteria (Marty *et al.*, 2002) but can also be present in other phytoplankton types (Jeffrey *et al.*, 2011). Station 1 showed a different pattern. *Synechococcus* counts by flow cytometry were lowest in winter and highest in spring in each set of cruises. Zeaxanthin did not match this pattern at station 1 with summer being higher than spring samples (no data were available for winter for station 1).

Picoeukaryotes were not detected by flow cytometry in the spring cruises but were present at all stations in the winter cruises (Supplementary Table S6). In summer, picoeukaryotes were found in all stations in 2016, but only at station 4 in 2015. The concentrations of all detected pigments and of all flow cytometry data are given in Supplementary Table S4 and S6, respectively.
